# Chemical inactivation of a bacterial immune system de-domesticates a temperate phage and promotes its spread

**DOI:** 10.64898/2026.05.14.725270

**Authors:** Yanyao Cai, Jaka Jakin Lazar, Yun Shi, Biswa P. Mishra, Thomas Ve, Anna Dragoš, Joseph P. Gerdt

## Abstract

Microbial communities shape hosts and ecosystems, yet the behaviors of these microbes are themselves controlled by mobile genetic elements. These mobile genetic elements (MGEs) compete and influence one another’s distributions. For example, many anti-phage immune systems are carried by prophages, plasmids, and integrative and conjugative elements (ICEs). It is unclear how important each of these interference mechanisms is for the spread of different mobile elements. Chemical inhibitors of these mechanisms of inter-MGE competition can clarify their mechanisms and significance in diverse environments. We discovered chemical inhibitors of SpbK—an antiphage defense encoded by an ICE in several strains of *Bacillus subtilis*. It is believed to prevent the dissemination of SPβ -like temperate phages, forcing SPβ to remain ‘domesticated’ in its ICE-containing host. Chemical inhibition of SpbK dramatically improved the escape of infectious SPβ phages capable of disseminating and forming new lysogens. Furthermore, SpbK is part of a widespread class of Toll/interleukin-1 receptor (TIR)-domain containing enzymes that are prevalent both in prokaryotic antiphage immune systems and eukaryotic immune signaling. Our inhibitors reveal two distinct mechanisms of inhibiting TIR-domain enzymes. These models of inhibition may broadly apply across immune proteins, enabling inhibition of many TIR immune systems that influence phage spread as well as plant immunity and animal cellular signaling.

## INTRODUCTION

Microbial communities play central roles in shaping host biology and ecosystem function^1–3^, yet their behaviors are profoundly influenced by the mobile genetic elements (MGEs) they harbor^4,5^. MGEs (e.g., plasmids, bacteriophages, and integrative and conjugative elements (ICEs)) can either integrate into the genomes of their host bacteria or remain as episomal elements. In either case, they both vertically transmit by replicating and passing onto their host cell’s progeny, as well as horizontally transmit between cells in the same physical environments^4^. Importantly for their hosts and microbial communities, these MGEs can add and disrupt genes that change the behaviors of their hosts. For example, MGEs frequently confer antibiotic resistance, virulence, and metabolic pathways to their hosts, which can dramatically influence host physiology^6^, microbial community composition^7,8^, and microbe-host symbioses^9^.

MGEs do not interact with bacteria in isolation. Rather, genetic elements engage in complex competitive and cooperative interactions with each other that shape their persistence and distribution across microbial populations. MGEs can improve the fitness of their host as well as other MGEs in that host^10,11^. They can conversely impose a fitness cost on the host and other co-resident MGEs^12,13^. Perhaps most dramatically, a prophage kills its host (along with all its MGEs) when it converts to its lytic cycle to transmit and colonize new hosts^14^. To avoid this costly outcome, MGEs have acquired an arsenal of anti-phage immune systems to presumably ‘domesticate’ co-resident prophages, preventing them from lysing the host as well as preventing lysis by newly infecting lytic phages^15,16^. We are gaining an increased appreciation for the abundance, distribution, and diversity of these inter-MGE inference genes. However, their biochemical mechanisms and their relative contributions to controlling the spread of different MGEs remain nascently understood.

Small-molecule inhibitors provide a powerful approach to dissect these competitive mechanisms and to evaluate their ecological and evolutionary significance in complex environments. A couple of previous reports have established the feasibility of immune system inhibitors to sensitize bacteria to lytic phages^17,18^, which encourages optimism for the viability of similar inhibitors to regulate MGE transfer as well. In this initial foray into controlling inter-MGE interactions, we chose to study the interaction between ICE*Bs*1 (an integrative conjugative element in *Bacillus subtilis*) and SPβ, a common *B. subtilis* prophage that confers bacteriocin production, spore resilience, and resistance to other phages, as well as possibly new metabolic pathways and antibiotic resistance to its hosts^19^. Specifically, we focused on the recently discovered protein SpbK encoded by ICE*Bs*1. This immune protein is reported to prevent SPβ from completing its lytic cycle, essentially domesticating the prophage and retaining it safely within hosts that harbor ICE*Bs*1^19–21^.

SpbK is part of a broader set of TIR (Toll/interleukin-1 receptor) domain-containing bacterial immune systems^20–22^. As with their human TIR homolog SARM1 (sterile alpha and TIR motif-containing 1)^23,24^, the TIR domains in both bacterial and plant immune systems generally possess self-association-dependent NADase activity that either depletes intracellular NAD⁺ (directly impacting cell viability)^25–28^ or modifies NAD⁺ into a series of signaling molecules including several non-canonical isomers of cADPR (cyclic adenosine diphosphate ribose) including 2′-cADPR^29–31^, 3′-cADPR ^30–32^ and N7-cADPR^33^, as well as His-ADPR (histidine-ADPR)^34^, ADPR-ATP^35^, pRib-AMP/ADP (phosphoribose-AMP/ADP)^35,36^ and di-ADPR^35^, which all serve to activate immune effectors. SpbK hydrolyzes NAD^+^ into N1-cADPR and ADPR after activation by YonE, a phage-encoded protein that is expressed either when the SPβ lysogen is activated to its lytic phase or if a YonE-containing lytic phage infects the cells and starts replicating (Fig. 1a)^20,21^. We aimed to develop chemical inhibitors of SpbK to probe its mechanism and significance for competition between mobile genetic elements and the dissemination of the SPβ lysogen.

**Figure 1.**
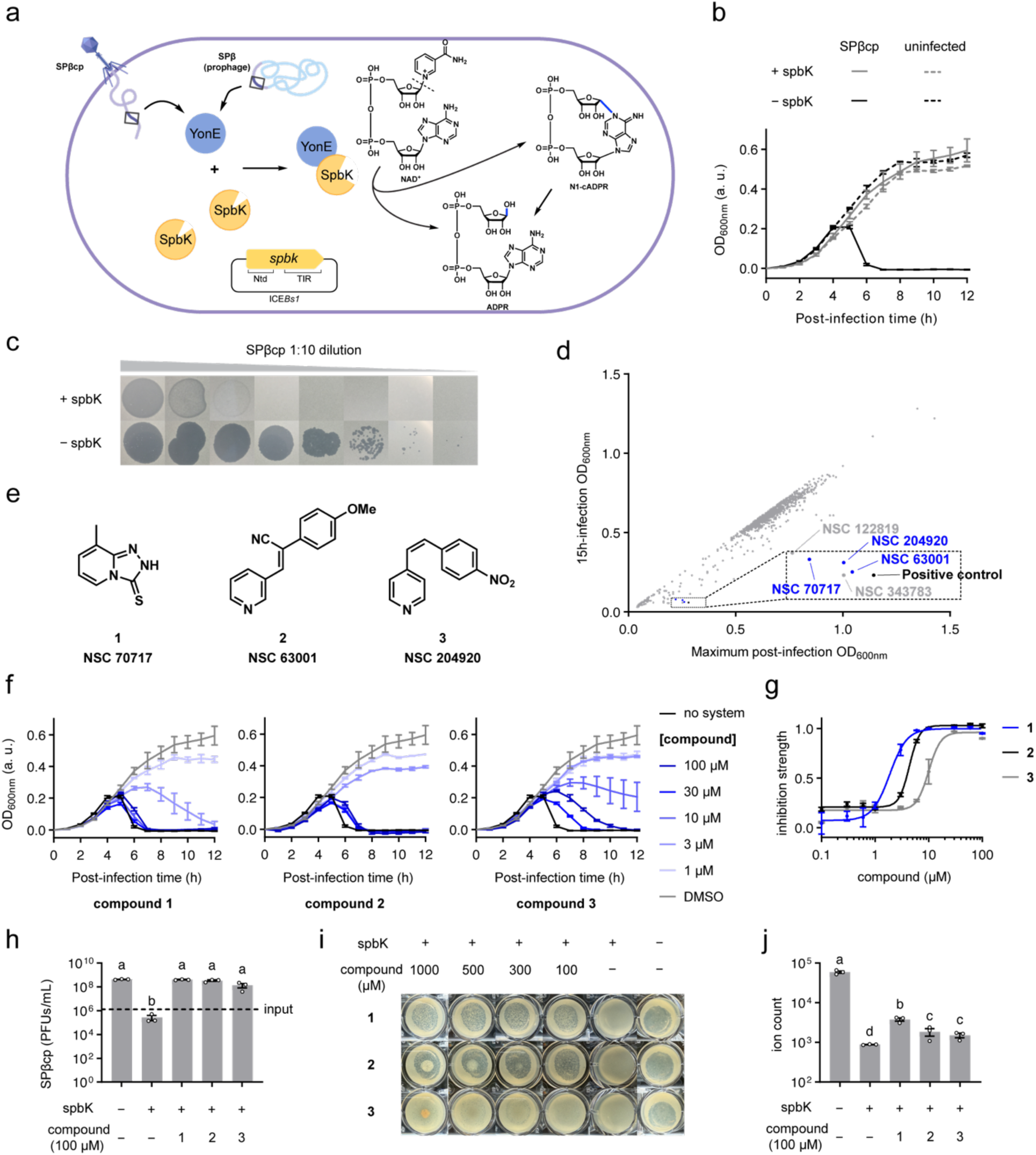
High-throughput screen identifies three small molecule inhibitors of the SpbK anti-phage defense. (a) Domain composition of *spbK* (Ntd: N-terminal domain, TIR: Toll/interleukin-1 receptor domain) on ICE*Bs1* and cartoon illustrating the immune mechanism of the SpbK system, which converts NAD^+^ into ADPR via hydrolysis or N1-cADPR (ultimately into ADPR) after activation by YonE expressed from an activated SPβ prophage or related obligately lytic phages like SPβcp. (b) Lysis curves of +SpbK strain of *Bacillus subtilis* (CMJ82, which is CU1050 with *spbK* inserted at the *amyE* locus) or –SpbK strain of *B. subtilis* (CU1050, a 168 mutant lacking its endogenous SPβ prophage and *spbK*-contained ICE*Bs1* locus) infected with SPβcp phage at multiplicity of infection (MOI) of 0.3. (c) Plaque images resulting from ten-fold serial dilutions of SPβcp phage spotted onto +SpbK strain (1st row) and –SpbK strain (2nd row). (d) Results from screening the NCI Diversity Set VII library, presented as a two-dimensional plot revealing potential inhibitors of the SpbK system, which allow initial bacterial growth to OD600nm ∼0.3, followed by lysis to a minimal OD600. The dashed box highlights compounds that provide lysis curves similar to the ‘Positive control’ (phage-infected –SpbK cells, black spot) ± 0.08 OD600nm units in the x-dimension and ± 0.03 OD600nm units in the y-dimension. (e) Chemical structures of compounds 1, 2 and 3 with corresponding NSC numbers. (f) Lysis curve of +SpbK strain treated with SPβcp phage (MOI = 0.3) and different concentrations of compounds 1, 2 and 3 compared to a control, –SpbK strain (black). (g) Dose-response curves of compounds 1, 2 and 3 measured with +SpbK strain treated with SPβcp phage (MOI = 0.3). See also Fig. S1. (h) SPβcp phage reproduction, quantified by measuring the plaque forming units (PFUs) 18 hours post-infection in the presence of compounds 1, 2, and 3 (100 µM). Input indicates the initial PFUs in the culture. ANOVA analysis of log10-transformed data is indicated by compact letter display. Bars with shared letters are not significantly different. (i) Plaque images resulting from SPβcp phage (∼10^4^ PFUs) spotted onto +SpbK strain with different concentrations of compounds 1, 2 and 3, compared to a control, –SpbK strain (last column). See also Fig. S2. (j) Intracellular NAD^+^ level (ion count represented by peak area under extracted ion chromatogram (EIC) of [NAD^+^–2H]^–^ (m/z 662.0887 – 662.1153)) measured by HPLC-HRMS from lysates extracted from +SpbK strain or control (–SpbK strain) under SPβcp phage infection at MOI ∼ 10 in the presence of compounds 1, 2, and 3 (100 µM each). ANOVA analysis of log10-transformed data is indicated by compact letter display. Bars with shared letters are not significantly different. In panels b, f-h, and j, data are represented as the mean ± SEM of n = 3 independent biological replicates. In panels h and j, individual replicates are displayed as circles.

We found multiple SpbK inhibitors that surprisingly exhibited at least two distinct mechanisms of inhibition. Our most potent inhibitor dramatically improved the release of transmissible SPβ, revealing the significance of this gene for inter-MGE competition. Remarkably, we found a natural product (nicotinamide) also inhibits this behavior, suggesting that metabolic environments play a role in this inter-MGE competition. Furthermore, SpbK belongs to a widely distributed family of Toll/interleukin-1 receptor (TIR)-domain-containing enzymes^22^, which are found in many bacterial antiphage systems as well as eukaryotic immune signaling pathways^37,38^. Given that multiple relatively simple small molecule inhibitors of SpbK were discovered from small screening libraries, we are optimistic that chemical probes may be developed to control many TIR-domain enzymes to study and control the spread of MGEs in bacterial communities and immune signaling in macroscopic eukaryotes.

## RESULTS

### High-throughput screen identified small molecules that disrupt the SpbK antiphage immune process

We first devised a screen to discover chemical inhibitors of the SpbK immune protein of ICE*Bs*1. Consistent with previous reports^20^, the SpbK system protected *B. subtilis* populations from complete lysis by an obligately lytic “clear plaque” mutant of SPβ (SPβcp) phage in liquid culture (Fig. 1b). Furthermore, on solid media, the SpbK system conferred greater than ten-thousandfold protection against phage infection. This was evidenced by the formation of clear SPβcp plaques on control cells lacking the system, whereas plaque formation was abolished in SpbK-expressing cells (Fig. 1c).

If a chemical could disrupt the SpbK immune process, then the SPβcp phage would resume lysis of the bacterial population. In liquid culture, the OD_600nm_ would initially increase and then drop at later time points as the culture lyses after 4 hours. Notably, this lysis process would appear differently than growth inhibition caused by any chemicals that are generally toxic to the bacteria. In that case, the bacteria would exhibit a low OD_600nm_ over the entire time course.

We leveraged this time-course OD_600nm_ measurement to screen the NCI Diversity Set VII library (1,581 compounds) for molecules that inhibit the SpbK immune system and therefore promote phage-induced lysis. Six compounds appeared to disrupt antiphage activity of SpbK resulting in a lysis curve resembling the no-system positive control (i.e., an early OD_600nm_ peak, followed by a decrease to a minimal OD_600nm_, reflected as a spot close to the positive control in the screening result plot [Fig. 1d]). Upon further investigation, three candidates (Fig. 1e) exhibited reproducible dose-dependent and phage-dependent host lysis (Fig. 1f-g, S1). In addition to promoting phage-induced cell lysis, an SpbK inhibitor should enable greater phage replication on SpbK-expressing hosts. Indeed, compounds 1, 2, and 3 all recovered the reproduction efficiency of the SPβcp phage on an SpbK-expressing *B. subtilis* host (Fig. 1h).

Given that some applications would require high inhibitor solubility and diffusion through semi-solid media/tissue, we evaluated the inhibitory activity of each compound on solid agar cultures of SpbK-expressing cells. Treatment with either compound 1 or 2 significantly promoted SPβcp phage infection, resulting in the formation of clear plaques comparable in size to the control (Fig. 1i). This lysis was phage-dependent, as exclusion of phage caused no impact on bacterial growth (Fig. S2a). In contrast, treatment with compound 3 yielded only sporadic clear plaques, indicating weaker efficacy. Compound 3 also precipitated out of the aqueous agar media at high concentrations. Therefore, the higher potency and solubility/diffusion of compounds 1 and 2 make them more promising for application to inhibit bacterial immunity in native environments (Fig. S2b-c).

### Intracellular NAD^+^ analysis suggests inhibition of SpbK NADase activity

To verify that the phage-promoting activity of these three inhibitors stems from the suppression of the SpbK immune system, we monitored their ability to prevent phage-induced SpbK-mediated depletion of intracellular NAD^+^. Infection with SPβcp at a high multiplicity of infection (MOI ∼10) ensured synchronous infection across the cell population to activate SpbK-mediated NAD^+^ cleavage. Analysis of the infected cell lysates via high-performance liquid chromatography-high resolution mass spectrometry (HPLC-HRMS) revealed that compound 1 significantly mitigated NAD^+^ depletion, although levels remained below those of uninfected controls (Fig. 1j). Compounds 2 and 3 also exhibited the capacity to partially preserve the NAD^+^ pool, but with significantly weaker efficacy than compound 1. Furthermore, treatment with compound 1 or 2 restored a normal high-MOI lysis curve, closely resembling the lysis kinetics of control cells lacking the SpbK system (Fig. S3). On the other hand, compound 3 produced an incomplete lysis curve within the same time interval, which is likely related to its weaker potency. Together, these data confirm that these compounds promote phage infection via disrupting the SpbK immune system in *B. subtilis*.

### Checkerboard analysis reveals a weak additive interaction between the SpbK inhibitors

We first classified these inhibitors into two categories based on their structural characteristics: inhibitors 2 and 3 carry an exposed pyridine ring, whereas inhibitor 1 contains a distinct heteroaromatic ring, 1H-1,2,4-triazole-3-thione. Given their structural differences, we hypothesized that they might inhibit SpbK via different mechanisms. If so, they could exhibit synergy and provide a more effective inhibitory cocktail. To test this hypothesis, we performed a synergy checkerboard assay with the potent and structurally distinct SpbK immune system inhibitors 1 and 2. Instead of synergy, the combination demonstrated only a weak additive effect, as highlighted by the white circle (fractional inhibitory concentration index (FICI) = 0.5–1.0) in the assay heatmap (Fig. 2a, S4).

**Figure 2.**
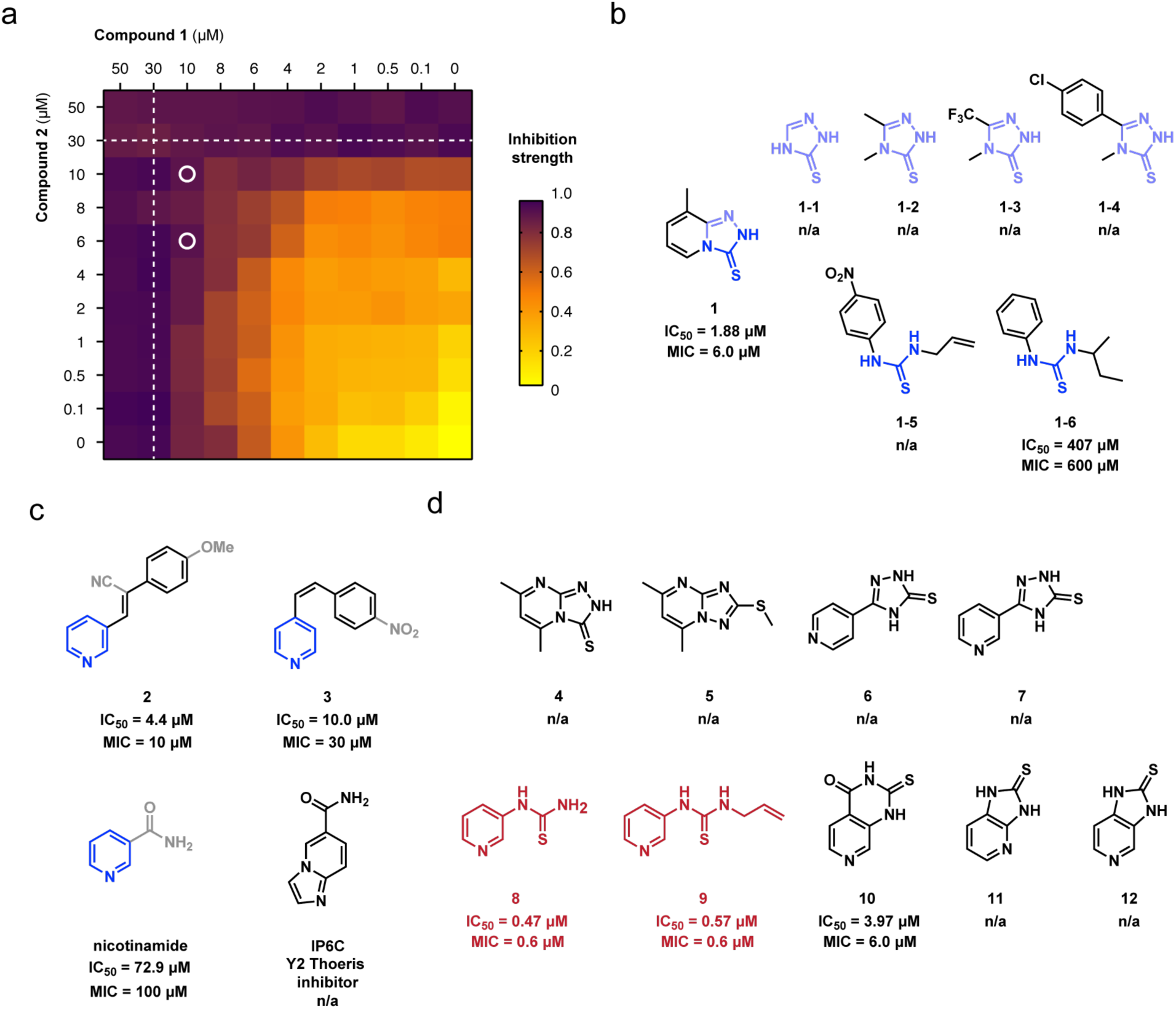
Interaction between inhibitors and structure-activity relationship study reveals optimal SpbK inhibitors. (a) Heatmap representing a synergy checkerboard analysis indicates the weak additive effect of compounds 1 and 2 inhibiting the SpbK immune system under SPβcp phage infection (MOI = 0.3). Inhibition strength values represent means of 3 independent biological replicates. White dashed lines indicate the minimal inhibitory concentration (MIC) of compound 1 or 2 alone (MIC here defined as inhibition strength > 0.9). White circles indicate combinations exhibiting a fractional inhibitory concentration index (FICI) between 0.5 and 1.0. No FICI values were <0.5, which would indicate synergy. See also Fig. S4. (b) Chemical structures and IC_50_ and MIC values of compound 1 and analogs 1-1 to 1-6. See also Fig. S5 and S6. (c) Chemical structures and IC_50_ and MIC values of compounds 2, 3, nicotinamide, and IP6C. See also Fig. S6 for nicotinamide and S5 for IP6C. (d) Chemical structures and IC_50_ and MIC values of compounds 4–12. See also Fig. S5 and S7. In panels b and c, the tested maximum concentration of compounds is 1 mM. In panel d, the tested maximum concentration of compounds is 100 µM. “n/a” means no activity.

### Structure-activity relationship study guides the discovery of inhibitors with improved potency

To explore the structural features required for each inhibition mode and further identify more potent compounds, we conducted a structure-activity relationship (SAR) study on our initial hits: inhibitors 1, 2, and 3. The most potent *in cellulo* inhibitor, compound 1, attracted our attention first. As heteroatoms frequently play an essential role in mediating protein-ligand interactions^39^, we focused on the heteroatom-rich core moiety of inhibitor 1, particularly retaining the thiourea, which is a privileged moiety capable of hydrogen bonding, electrostatic interactions, and metal coordination^40,41^. Based on this scaffold, we evaluated a series of analogs, including derivatives bearing trifluoromethyl and *p*-chlorophenyl groups (compounds 1-2 to 1-4), as well as simplified thiourea derivatives (compounds 1-5 and 1-6). Compounds 1-1, 1-2, 1-3, and 1-4 were all inactive, indicating that the second fused aromatic ring of inhibitor 1 is critical for activity (Fig. 2b, S5). Another thiourea- and aromatic ring-containing analog (compound 1-6) also inhibited SpbK (Fig. 2b, S6), providing affirmation for the value of having both a thiourea and an aromatic moiety.

We next investigated the second class of inhibitors, compounds 2 and 3, which notably share a common 4-styrylpyridine scaffold. Pyridine-containing inhibitors of the human TIR-domain NADase SARM1 have recently been shown to form covalent conjugates with ADPR via TIR-domain-catalyzed base exchange with NAD^+42–44^. Therefore, we hypothesize that compounds 2 and 3 inhibit SpbK-mediated NAD^+^ depletion through a similar mechanism: by directly forming conjugates with ADPR in the enzyme active site. The styryl group appears to improve *in cellulo* potency, as the related natural metabolite nicotinamide required higher concentrations to inhibit SpbK (Fig. 2c, S6). Intriguingly, not every base-exchangeable heterocycle inhibits the antiviral activity of SpbK. Specifically, we tested IP6C, a molecule that is readily linked to ADPR by SpbK ^18,22^, but it failed to inhibit SpbK-mediated antiphage activity (Fig. 2c, S5). Therefore, although multiple heterocycles can be base-exchange substrates of SpbK, a pyridine ring appears to be preferred for inhibition of SpbK-induced abortive infection.

Guided by our understanding of the traits driving the bioactivity of both inhibitor classes, we attempted to merge these features to generate a hybrid molecule with the goal of developing a more potent SpbK inhibitor. First, we introduced a nitrogen atom into the aromatic ring of inhibitor 1 to generate a pyridine-like moiety (compounds 4 and 5), a feature critical for base exchange type inhibition. However, this modification resulted in a complete loss of bioactivity (Fig. 2d, S5). Directly connecting a pyridine ring to a 1H-1,2,4-triazole-3-thione core also yielded inactive compounds 6 and 7 (Fig. 2d, S5). Finally, we tested a series of thiourea derivatives containing a pyridine ring. This straightforward strategy yielded our most potent SpbK inhibitor, compound 8, which exhibited a minimal inhibitory concentration (MIC) tenfold lower than that of inhibitor 1 (Fig. 2d, S7). The introduction of an allyl group to the thiourea of 8 had no effect (compound 9 [Fig 2d, S7]). Subsequent structural constraints were tested. Introducing a ketone to fix the thiourea within a ring resulted in poor aqueous solubility and reduced potency (compound 10 [Fig 2d, S7]). Furthermore, linking both thiourea nitrogen atoms to the pyridine ring completely abolished activity (compounds 11 and 12 [Fig 2d, S5]). Therefore, the most potent inhibitors (compounds 8 and 9) contain both a thiourea (like compound 1) and a pyridine (like compounds 2 and 3); however, they do not restrict the thiourea into a ring like in compound 1.

### *In vitro* NADase activity assay revealed two distinct inhibitory pathways against SpbK immunity

As mentioned above, based on structural characteristics, we hypothesized that compounds 2, 3, and 8 would form covalent conjugates with ADPR like pyridine-containing inhibitors of SARM1 (Fig. 3a), but compound 1-6 would not. Although compound 1 has no pyridine, it does have a nucleophilic amine that might be capable of displacing nicotinamide as well. To test for base-exchange reactions generating these possible conjugates, we expressed and purified SpbK as a maltose-binding protein fusion (MBP-SpbK) along with its phage-derived activator, YonE. We incubated MBP-SpbK/YonE with NAD^+^ under the treatment of each inhibitor and used high performance liquid chromatography coupled to high resolution mass spectrometry (HPLC-HRMS) to detect the formation of conjugates, as well as monitor the depletion of NAD^+^ and formation of N1-cADPR and ADPR. We also included compound 1-5 (analog of 1-6) in this *in vitro* analysis in case it inhibits SpbK but fails to access the protein *in cellulo*. Indeed, all inhibitors delayed the production of N1-cADPR and ADPR by SpbK (including even compound 1-5, Fig. 3b). Furthermore, we detected base-exchange products of the expected m/z between NAD^+^ and all four potential base-exchanging inhibitors, including compound 1 (Fig. 3c). The identity of these inhibitor-ADPR conjugates was further validated via tandem mass spectrometry (MS/MS), which revealed characteristic ADPR fragmentation patterns for each of the conjugates (Fig. 3d, S8a-b). The formation of these inhibitor-ADPR conjugates was further supported by *in situ* ^1^H-NMR monitoring of the reaction. Specifically, transglycosidation was evidenced by the emergence of new resonances in the anomeric region (5.5 to 6.0 ppm), indicating the covalent attachment of the ribose anomeric carbon to the inhibitors (Fig. 3e and S9). As hypothesized from its lack of a heterocyclic amine, compound 1-6 and its analog compound 1-5 did not form conjugates with ADPR detectable by HPLC-HRMS or ^1^H-NMR (Fig. S8c, S9). Therefore, although compounds 1, 2, 3, and 8 likely inhibit SpbK via the formation of ADPR conjugates in the enzyme active site, compound 1-6 and its analog compound 1-5 inhibit SpbK via different pathway—either by binding the active site differently or via a separate route.

**Figure 3.**
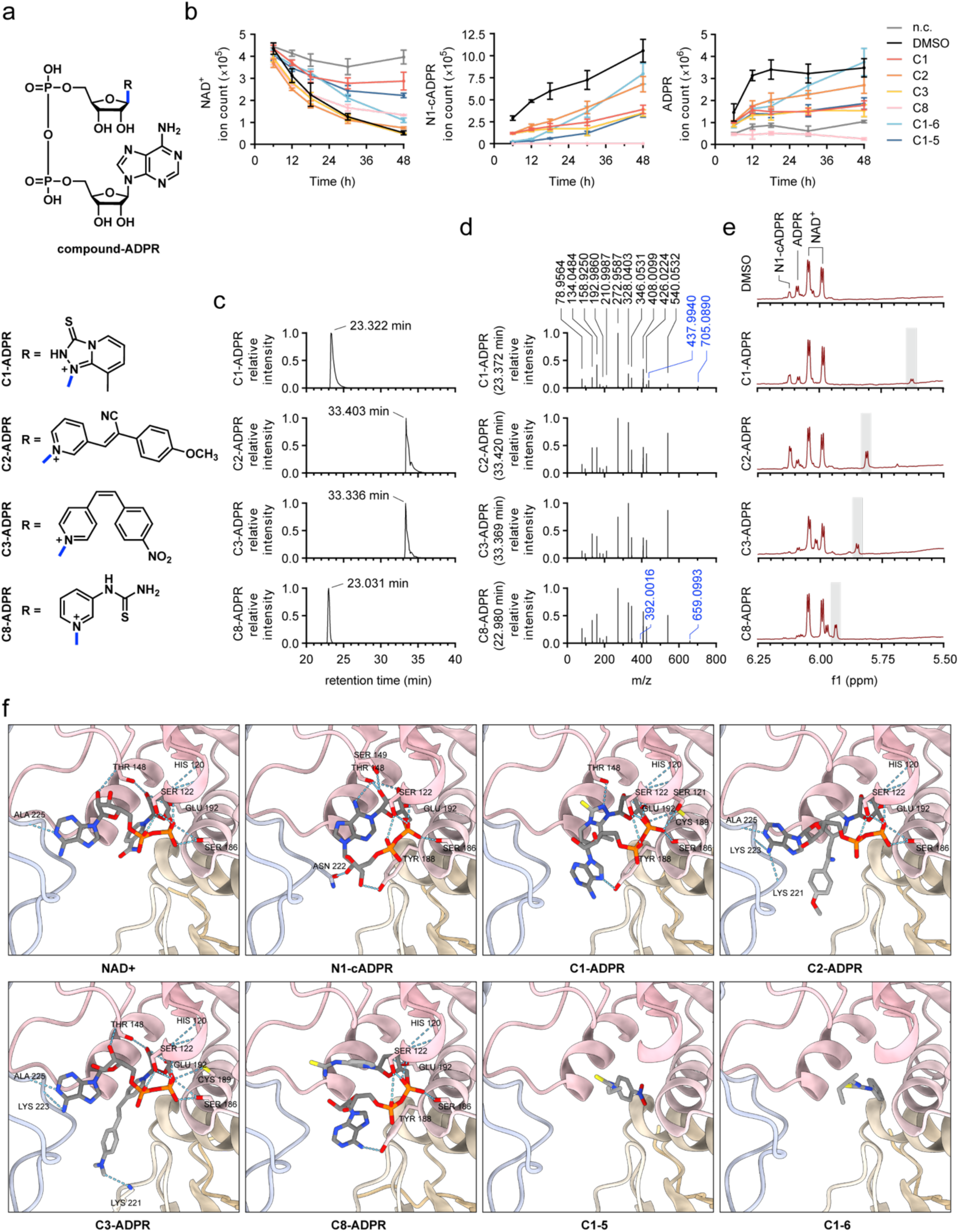
*In vitro* analysis of SpbK-mediated NAD^+^ depletion reveals conjugate-forming and non-conjugate-forming mechanisms of inhibition (a) Chemical structures of compound-ADPR conjugates formed from the SpbK-catalyzed reaction between NAD^+^ and compounds 1, 2, 3, or 8. (b) HPLC-HRMS-based time-course kinetics of NAD^+^, N1-cADPR, and ADPR during the reaction (500 µM NAD^+^, 2 µM MBP-SpbK, 30 µM YonE and 5 mM inhibitor or DMSO as control in 50 mM HEPES, 150 mM NaCl in 95% D_2_O) at 30 °C. “n.c.” means no SpbK enzyme was added. Ion count was quantified by integrating the peak area in extracted ion chromatograms (EICs) for m/z 540.0431-540.0649 (matching N1-cADPR [M–H]^–^), m/z 662.0887-662.1153 (matching NAD^+^ [M–2H]^–^) and m/z 558.0532-558.0756 (matching ADPR [M-H]^-^). Data are represented as the mean ± SEM of n = 3 independent biological replicates. (c) Extracted ion chromatograms (EICs) for m/z 705.0758-705.1040 (matching C1-ADPR [M-2H]^-^), 776.1333-776.1643 (matching C2-ADPR [M-2H]^-^), 766.1128-766.1434 (matching C3-ADPR [M-2H]^-^) and 693.0760-693.1038 (matching C8-ADPR [M-2H]^-^) from reaction component analysis after 18 hours in Fig. 3b. Peak intensities were normalized by the strongest peak in each chromatogram. (d) MS/MS spectra of corresponding compound-ADPR conjugates in Fig. 3c. Peak intensities were normalized by the strongest peak in each spectrum. The shared characteristic m/z’s generated from the ADPR moiety of the conjugates are labeled in black. Other characteristic m/z’s are labeled in blue. See Fig. S8 for predicted fragment structures. (e) ^1^H-NMR spectra (5.50-6.25 ppm) of each reaction mixture at 12 hours in Fig. 3b. The signal of the proton connected to the ribose anomeric carbon in each compound-ADPR conjugate is highlighted with a grey shadow. See Fig. S9 for full spectra. (f) Enlarged cutaway views of the active-site region (located at the interface between TIR subunits within each strand) in the Boltz-2 co-folded complexes between ligands (NAD⁺, N1-cADPR, C1-5, C1-6 and C1-, C2-, C3-, C8-ADPR conjugates) and a SpbK TIR domain (residues: 113-266) hexamer from the SpbK TIR filament cryo-EM structure (PDB: 9PHB). SpbK residues predicted to mediate binding of NAD⁺, N1-cADPR, and ADPR conjugates are indicated, with hydrogen bonds indicated as light blue dashed lines.

Further kinetic analysis of the reaction components (NAD^+^, N1-cADPR, and ADPR) revealed distinct differences among the inhibitors (Fig. 3b). Compound 1 matched our expectations for a ‘prodrug’ that forms the C1-ADPR conjugate, which ultimately inhibits SpbK (analogous to inhibitors of SARM1). Specifically, NAD^+^ was depleted at initial time points during the formation of C1-ADPR, but NAD^+^ depletion was halted at later time points after C1-ADPR was formed.

Surprisingly, inhibitors C2 and C3 failed to slow NAD^+^ depletion under these experimental conditions. Likewise, they were significantly worse than C1 at inhibiting NAD^+^ depletion *in cellulo* (Fig. 1j). However, both compounds significantly inhibited the formation of N1-cADPR and ADPR by SpbK. Therefore, we conclude that these inhibitors depleted NAD^+^ but diverted flux away from N1-cADPR and ADPR by instead generating C2- and C3-ADPR conjugates. In contrast to C1, the C2- and C3-ADPR conjugates failed to inhibit further depletion of NAD^+^. We propose two models to explain this observation. First, these conjugates may simply inhibit NAD^+^ depletion less strongly than C1-ADPR, and the *in vitro* experiment conditions fail to reveal this slight inhibition that may still occur in true cellular conditions. The weaker inhibition may be caused by weaker binding of the ADPR conjugates to SpbK; however, computational modeling of all three ADPR conjugates into SpbK by Boltz-2 did not suggest tighter binding by C1-ADPR (Fig. 3f, Table S1). Instead of weaker binding, the C2- and C3-ADPR conjugates may also be produced slower or depleted faster by SpbK, thereby keeping their concentrations lower and less able to inhibit SpbK. As an alternative model, C2- and C3-ADPR may inhibit the abortive infection of SpbK through a mechanism that does not require maintenance of NAD^+^. Specifically, these conjugates may ‘replace’ the function of NAD^+^ to prevent abortive infection—at least for long enough to allow replication of the phage. Although C2- and C3-ADPR have never been tested in this context, several other base-exchanged analogs of NAD^+^ have been shown to substitute for NAD^+^ as substrates in enzyme-catalyzed reactions.^45–49^ We note that the exact cellular mechanism by which NAD^+^ depletion induces abortive infection is yet unclear. Therefore, it is plausible that C2-ADPR and C3-ADPR may delay SpbK-mediated abortive infection by mimicking NAD^+^ instead of preventing its depletion. These two “competing” models are not mutually exclusive—an inhibitor-ADPR conjugate may *both* inhibit SpbK and mimic NAD^+^ in the yet-unknown NAD^+^-dependent process(es) that prevent abortive infection.

Since the *in cellulo*-active, exocyclic thiourea-containing compound 1-6 and its analog (compound 1-5) are incapable of forming ADPR conjugates, we hypothesized that they would simply prevent SpbK-catalyzed NAD^+^ cleavage. Indeed, they slowed NAD^+^ depletion *in vitro* (Fig. 3b). These compounds also exhibited a greater ability to prevent N1-cADPR formation at early time points compared to the pure base-exchange inhibitors (compounds 1, 2 and 3). This finding suggests that the exocyclic thiourea moiety uniquely prevents the formation of N1-cADPR.

As expected, one of our most potent inhibitors, compound 8, exhibited the strongest *in vitro* inhibition of N1-cADPR generation. This compound combines both aforementioned structural features: a pyridine ring to participate in the base-exchange reaction, and an exocyclic thiourea to halt N1-cADPR formation. Its remarkable potency was evidenced by the complete absence of detectable N1-cADPR throughout the entire reaction process, as measured by HPLC-HRMS. Like the pyridine-containing base-exchange inhibitors 2 and 3, it also allowed substantial NAD^+^ depletion in this assay, further confirming that complete inhibition of NAD^+^ depletion is not necessary to prevent abortive infection by SpbK.

Together, our results support categorizing the inhibitors based on the earlier noted structural features: (1) presence of a nucleophilic heterocyclic amine for conjugation to ADPR and (2) a thiourea moiety, which appears to promote SpbK inhibition. Compounds 1 and 8 have both features, albeit the endocyclic thiourea of compound 1 may not enable the full inhibition of N1-cADPR production. Compounds 2 and 3 are base-exchangers that lack the thiourea. Compounds 1-5 and 1-6 have the thiourea but lack base-exchange ability. In summary, there are at least 2 distinct mechanisms of inhibition—one involving base-exchange and formation of covalent ADPR conjugates and another than does not. Even among inhibitors that form ADPR conjugates, there may be different ultimate mechanisms of delaying abortive infection.

### Compound 8 rescues lysogenic-to-lytic induction of native SPβ prophage

Now with the ability to inhibit SpbK, we sought to leverage the chemical inhibitors to regulate ICE*Bs*1-mediated control of its co-resident, native SPβ prophage in *B. subtilis*. This contrasts with our previous experiments, which used an obligately lytic mutant of the SPβ phage. That decision was out of experimental ease, which has also driven most other studies of antiphage immune systems to deploy lytic phages^51–53^. However, in many environments, temperate phages capable of lysogeny (like SPβ) are estimated to be far more abundant those that are obligately lytic^54,55^. These temperate phages can integrate into host genomes and lay dormant, passively replicating along with the host genome into each of the host’s progeny. Like other MGEs, these integrated prophages influence the genes and gene expression of their hosts. Unlike other MGEs, however, many such prophages convert to their lytic cycle at a low baseline rate to spread to infect new hosts^56,57^. This rate of spontaneous lytic ‘induction’ is also tuned through sensing diverse endogenous and environmental factors to increase spread to new hosts when the conditions favor it^57–59^. This induction kills the host and co-resident MGEs. It is this context in which SpbK is hypothesized to natively function—it arrests the productive activation of SPβ, preventing its horizontal spread^19–21^. Therefore, our inhibitors of SpbK provide the opportunity to control and study the ICE-encoded anti-phage defense within its native context of hypothetically regulating the spread of a co-resident SPβ prophage.

The well-studied *B. subtilis* strain 168 (ATCC 23857) natively harbors both the integrative and conjugative element ICE*Bs1* (which encodes SpbK) and the SPβ prophage. SPβ spontaneously activates to release low titers of infectious viral particles, and the rate of induction can be substantially increased by DNA damage via exposure to mitomycin C or UV radiation.^19^ However, in the presence of ICE*Bs*1, SpbK prevents most of these induced phages from completing their lytic cycle and forming infectious viruses that can spread to new hosts^20,21^. We hypothesized that SpbK inhibition would restore the release of infectious SPβ. Indeed, treatment with compound 8 rescued the replication efficiency of the SPβ phage (Fig. 4a). This improvement in phage release was evident for both mitomycin-induced phage activation and spontaneous phage activation. Importantly, compound 8 did not significantly promote SPβ release from a control natural isolate that is closely related to 168 but lacks ICE*Bs*1 (MB8_B7, Fig. 4b). This control demonstrates that the promotion of SPβ release by compound 8 was specifically due to SpbK inhibition. Therefore, compound 8 suppresses the native SpbK system *in cellulo*, promoting the spread of native co-resident SPβ. This result validates the importance of SpbK in regulating the transfer of SPβ (and likely related prophages). Furthermore, these inhibitors (and other current and future inhibitors of bacterial immune systems) are viable tools to study the influences of MGEs on the ability of temperate phages (as well as lytic phages) to spread and shape native microbial communities.

**Figure 4.**
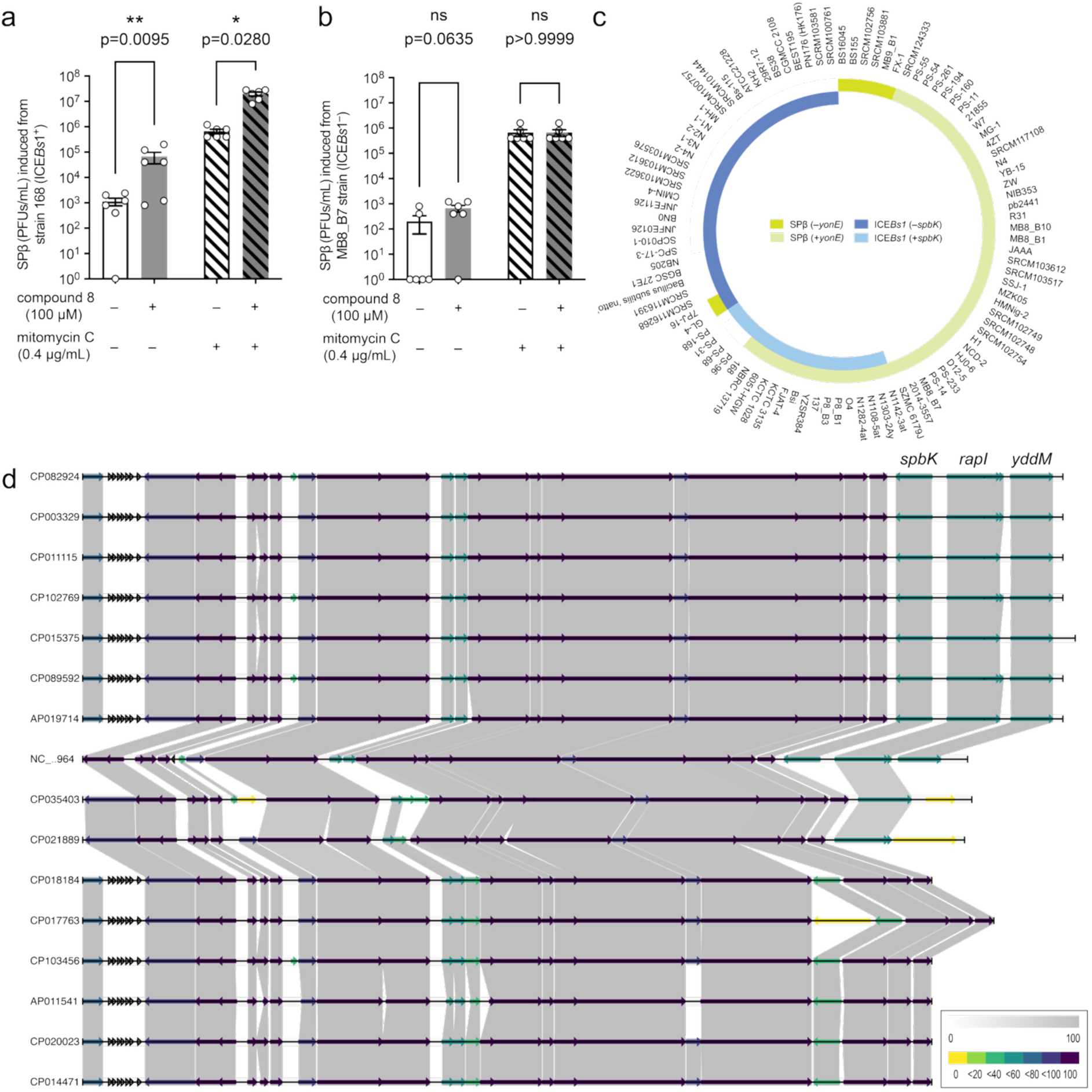
Impact of SpbK on SPβ prophage induction and co-existence of ICEBs1 and SPβ. (a–b) Level of infectious SPβ phage, quantified by counting PFUs released from (a) a +ICE*Bs1 B. subtilis* (strain 168) or (b) a –ICE*Bs1 B. subtilis* (strain MB8_B7) in the presence/absence of 100 µM compound 8, with and without induction by mitomycin C (0.4 µg/mL). Significance was determined by lognormal one-way ANOVA analysis with Tukey’s multiple comparisons test. Adjusted p values are displayed above comparisons between samples +/– compound 8. Data are represented as the mean ± SEM of n = 6 independent biological replicates, which are each displayed with a circle. (c) Graphical representation of the co-occurrence of ICE*Bs1* strains containing/lacking *spbK* with the SPβ prophage containing/lacking the SpbK-inducer *yonE* on each of 97 *B. subtilis* genomes. See also Table S2. Significant negative association between ICEBs1 and SPβ-like elements (p < 0.0001) and significant positive association between *spbK* and *yonE* (p = 0.021) was confirmed by a chi-square test. (d) Synteny plot generated by Synphage tool^50^ of the 16 representative ICE*Bs*1 sequences with some containing *spbK*. Color bar represents the percentage of conservation within the displayed sequences.

### *spbK* and SPβ frequently co-exist in *B. subtilis* genomes

Given the dramatic impact of SpbK on the potential for SPβ to spread, we sought to further understand the widespread significance of the SpbK immune system and its evolutionary linkage with SPβ lysogeny. We analyzed 97 diverse *B. subtilis* genomes that carried ICE*Bs1*, an SPβ-like prophage, or both. We then asked if ICE*Bs*1 and SPβ frequently reside in the same host and if the presence of ICE*Bs*1-encoded *spbK* and its phage-encoded activator *yonE* influence their stable co-existence. Since *spbK* may protect the ICE*Bs*1 (and the host) from lysis by SPβ, we hypothesize that *spbK* may be maintained to a higher degree in genomes harboring SPβ-like lysogens. First, we found that most strains either carried only ICE*Bs*1 (35/97) or only SPβ (44/97) but not both elements (18/97, ∼19%) (Fig. 4c, Table S2). However, of the 18 strains where ICE*Bs*1 and SPβ co-existed, nearly all possessed *spbK* within the ICE*Bs*1 and *yonE* within the SPβ (16/18) (Fig. 4c, Table S2).

Although *yonE* is present in the majority of observed SPβ-like lysogens (53/62, ∼85%), *spbK* is in a sizable minority of ICE*Bs*1-positive genomes (20/53; ∼38%). The *spbK* gene resides in a terminal region of the ICE*Bs1*, along with adjacent genes *rapI* and *yddM*. This terminal region is sometimes entirely absent from ICE*Bs1*, and sometimes it is present but lacks the *spbK* gene (Fig. 4d). Statistical analysis revealed that the presence of *spbK* within ICE*Bs1* is significantly linked to presence of *yonE* within SPβ (p = 0.021), but not to the SPβ prophage in general (p = 0.156). Therefore, *spbK* may be lost from ICE*Bs*1 when *yonE*-containing SPβ lysogens are not present to necessitate abortive infection protection from prophage induction.

Overall, this pattern of *spbK* and *yonE* presence across 97 *B. subtilis* isolates collected worldwide suggests that stable coexistence of ICE*Bs*1 and SPβ depends on the presence of *spbK* and its phage-expressed partner *yonE*. Acquisition and/or maintenance of the *spbK*-encoded abortive infection system is linked to lysogeny by SPβ carrying an element (*yonE*) vulnerable to this system. The inhibitors in this work may be employed to directly test the significance of SpbK for SPβ, ICE*Bs*1, and their host bacteria in native environments. Naturally occurring pyridine-containing metabolites like nicotinamide may also already be reducing the protective effect of *spbK* in some environments, thereby decreasing the selective pressure for ICE*Bs1* to acquire or maintain this immunity gene and ultimately changing the interplay between mobile genetic elements within *B. subtilis*.

## DISCUSSION

We discovered a series of compounds that disrupt the antiviral immunity conferred by SpbK. These inhibitors fall into two distinct mechanistic classes—ones that are substrates of the enzyme and form covalent ADPR conjugates and those that do not. With sub-micromolar MICs, some of these compounds are the most potent inhibitors of anti-phage immunity reported to date^17,18,60^. These inhibitors promote the release of infectious SPβ phage particles from lysogenized hosts. Furthermore, the target immunity protein, SpbK, frequently co-occurs with *yonE*-containing prophages in natural isolates. Therefore, we can envision the usage of these inhibitors as tools to study and control phage-host dynamics in natural environments.

Notably, SpbK is hypothesized to regulate the transmission of temperate phages like SPβ. Due to their ease of study, lytic phages have primarily been used to examine the impact of bacterial immune systems on phage-host dynamics^51–53^. However, in nature, anti-phage immune systems likely combat phages capable of lysogeny, given the prevalence of temperate phages in many environments^54,55^. Therefore, SpbK and dozens of other immune systems likely regulate the lysis of host bacteria and transfer of genetic information by temperate phages. Future studies chemically inhibiting SpbK and other immune systems in native microbial communities are now feasible to test this hypothesis. Furthermore, these inhibitors may be deployed on their own or in conjunction with lysis-promoting compounds^59,61^ as technologies to promote phage-induced lysis of bacteria to optimize productive microbial communities or eradicate deleterious bacteria.

Beyond SpbK, TIR-domain containing enzymes are widespread in bacterial antiviral immune systems^30,37^ and in eukaryotic signaling across plants and animals^37,38^. The relative ease with which we found several inhibitors from a small screen of 1,581 compounds suggests that many TIR domain enzymes may be chemically inhibited with minimal upfront effort. Indeed, similar heterocyclic compounds have been shown to inhibit the human NADases SARM1 and CD38^42–44^, as well as the bacterial type II Thoeris^18^. In all these cases, the inhibitors are substrates of the enzyme and form ADPR conjugates. Therefore, directed screening of small libraries of heterocycle-containing (especially pyridine-containing) compounds may rapidly reveal inhibitors of wide panels of TIR-domain containing enzymes in both bacteria and eukaryotes. We note that non-conjugate-forming inhibitors were also found and may likewise be broadly fruitful, as this mechanism has been successful against human SARM1, as well^62^.

Our findings also reveal the promise of discovering both ‘broad spectrum’ and selective TIR inhibitors. These are each valuable for their own applications. A ‘broad-spectrum’ inhibitor will find widespread utility, and a selective inhibitor will provide confidence in probing the specific enzyme of interest. We have now found that nicotinamide inhibits both SpbK and type II Thoeris, whereas IP6C only inhibits Thoeris and compounds 1, 2, and 3 selectively inhibited SpbK^18^. Also, we recently found that compound 1-6 promiscuously inhibits the TIR-and SIR2-containing type I Thoeris immune system (unpublished). Therefore, we believe that a toolkit of TIR-domain inhibitors with different spectra of inhibition is well within reach.

This manuscript focused primarily on a library of synthetic compounds; however, we note that the natural metabolite nicotinamide also inhibits SpbK. Many other pyridine-containing metabolites also exist in nature, and several of these may prove effective conjugate-forming inhibitors of SpbK. Therefore, environmental metabolites could be tuning this competition between ICE*Bs*1-encoded SpbK and phages in nature. Namely, pyridine-rich environments may favor phage replication, whereas pyridine-poor environments would provide maximal SpbK protection to the ICE and the host. Furthermore, we predict that many other TIR domain-containing anti-phage defenses will likewise be inhibited by nicotinamide and related heterocycles in the environment—adding to the scope of metabolite-influenced phage-bacteria interactions^59,63,64^.

In conclusion, the SpbK protein that mediates bacteriophage competition with chromosomal elements in many strains of *B. subtilis* can now be chemically regulated by inhibitors with two distinct mechanisms. Since these inhibitors promote phage replication that both kills host bacteria and spreads temperate phages with their arsenal of host-modifying genes, they would be effective tools in studying and controlling the MGEs that shape microbial communities. In fact, natural heterocyclic metabolites may already be shaping phage-host dynamics in this way. Finally, our findings warrant optimism that chemical inhibitors are attainable for many other TIR-domain-containing enzymes in both prokaryotic anti-phage immunity and eukaryotic cell signaling.

## METHODS

### Bacteria strains and growth conditions

*B. subtilis* CU1050 was used as the host bacteria for major phage-infection-related experiments. This strain is a mutant of *B. subtilis* 168 and lacks the SPβ prophage and ICE*Bs1* (the element containing SpbK). *B. subtilis* 168 (+ICE*Bs1*) and *B. subtilis* MB8_B7 (–ICE*Bs1*) were used to evaluate the inhibition of SpbK in a native SPβ prophage-harboring background. *B. subtilis* CU1050 was used as the indicator strain for SPβcp titer. *B. subtilis* Δ6 was used as the indicator strain for SPβ titer. *E. coli* BL21(DE3) (New England Biolabs, C2527I) was used for protein expression. Either *E. coli* or *B. subtilis* strains were routinely grown in LB broth at 37 °C, 220 rpm. When appropriate, the media was supplemented with kanamycin (50 µg/mL) or ampicillin (100 µg/mL) to maintain the plasmids (MBP-SpbK carried by pET-28b, YonE carried by pMCSG7).

### Bacteriophage culturing and titer assay (SPβcp)

Bacteriophage SPβcp was generally cultured using the liquid culture method. 2 mL overnight culture of –SpbK *B. subtilis* (CU1050) was diluted 1:100 into 200 mL fresh LB in the sterile Erlenmeyer flask and incubated at 37 °C until OD_600nm_ achieved ∼0.2. Then 2 mL SPβcp stock solution (∼10^6^ PFUs/mL) was added, and the resultant culture was incubated at 30 °C, 220 rpm for 12-15 hours. After centrifugation at 10,000 × g for 10 min, the supernatant was sterilized by 0.22 µm filtration (Millipore Sigma, S2GPU02RE).

To titer SPβcp phage stocks, 0.2 mL overnight culture of –SpbK *B. subtilis* (CU1050) was mixed into 5 mL LB soft agar (0.5% w/v) at 55 °C and quickly poured on the top of 20 mL LB agar (1.5% w/v) in a Petri dish (100×15 mm, VWR, Cat#89038-968). 10 µL tenfold serially diluted SPβcp phage stock was dropped on the top agar. After the phage spots dried, the plate was incubated at 30 °C for 12-18 hours. Plaque-forming units (PFUs) were determined by counting the derived plaques.

### High-throughput screening assay

The NCI Diversity Set VII library of 1581 compounds was used for screening. The compounds from the library were prepared as 200 µM in 10 µL LB (4% DMSO) plated in 384-well plates (Corning, 3701), resulting in 50 µM final concentrations after addition of bacteria and phage. An overnight culture of +SpbK *B. subtilis* (CMJ82) was diluted 1:100 into fresh LB. Then 20 mL of this diluted *B. subtilis* culture was added to each well of the 384-well plates, followed by 1 hour incubation at 30 °C. After incubation, 10 mL of SPβcp phage (∼1.4 × 10^5^ PFUs/mL in LB) was added to each well. The plates were incubated at 30 °C in a Biospa8 (Biotek), and the OD_600nm_ in each well was recorded every 1 hour using a Synergy H1 plate reader (Biotek).

### Evaluation of SpbK inhibition in liquid media cultures

A 4 µL DMSO stock solution (100× final desired concentration) of each compound was mixed with 100 µL fresh LB in 1.7 mL sterile Eppendorf tube by vortexing, and 10 µL aliquot of each resulting solution was added into wells of 384-well plates (Corning, 3701). An overnight culture of +SpbK *B. subtilis* (CMJ82) was diluted 1:100 into fresh LB. Then 20 µL of diluted *B. subtilis* culture was added to each well of the 384-well plates, followed with 1 hour incubation at 30 °C. After incubation, 10 µL of SPβcp phage (∼1.4 × 10^6^ PFUs/mL in LB) was added to each well. (10 µL LB used in negative control.) Then the plate was incubated at 30 °C in a Biospa8 (Biotek), and the OD_600nm_ in each well was recorded every 1 hour using a Synergy H1 plate reader (Biotek).

The inhibition of SpbK under compound treatment and phage infection was calculated by

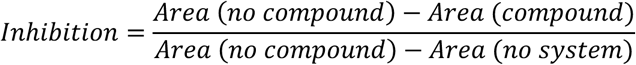

where “area” represents the integrated area under the bacterial lysis curve upon phage infection.

### SPβcp phage reproduction measurement in liquid media cultures

Phage reproduction was evaluated by quantifying the number of phages produced after infecting bacterial hosts. 10 µL of each inhibitor (400 µM in LB, 4% DMSO) was added into a well of a 384-well plate (10 µL LB used as negative control). An overnight culture of +SpbK *B. subtilis* (CMJ82) or –SpbK *B. subtilis* (CU1050) was diluted 1:100 into fresh LB. Then 20 µL of diluted *B. subtilis* culture was added to each well of 384-well plates, followed with 1 hour incubation at 30 °C. After incubation, 10 µL of SPβcp phage (∼1.4 × 10^6^ PFUs/mL in LB) was added to each well. The plates were incubated at 30 °C in a Biospa8 (Biotek) for 18 hours. After incubation, 10 µL infected culture was diluted with 1 mL fresh LB in a syringe, followed by passing through the 0.22 µm filter to remove the *B. subtilis*. The SPβcp concentration (PFUs/mL) was quantified on –SpbK *B. subtilis* (CU1050) using the small drop plaque assay described above.

### Evaluation of SpbK inhibition in solid media cultures

125 µL overnight culture of +SpbK *B. subtilis* (CMJ82) was mixed in 10 mL LB soft agar (0.5% w/v) at 55 °C. 250 µL of the resulting agar-bacteria mixture was quickly poured on the top of 1 mL LB agar (1.5% w/v) in each well of 24-well plate. Before the agar solidified, 2.5 µL of inhibitor (dissolved in DMSO at 100× final desired concentration) was immediately pipetted into the agar. After complete solidification of agar, 10 µL SPβcp phage (∼1.4 × 10^6^ PFUs/mL in LB, or fresh LB alone used as negative control) was dropped in the center of each well. After the spot dried in a biosafety cabinet, the plate was incubated at 30 °C for 18-24 hours.

### Evaluation of diffusion of inhibitors in solid media cultures

The protocol was modified based on the classic Kirby-Bauer disk diffusion susceptibility test protocol.^65^ 55 µL overnight culture of +SpbK *B. subtilis* (CMJ82) or –SpbK *B. subtilis* (CU1050) was mixed in 5 mL LB soft agar (0.5% w/v) at 55 °C and quickly poured on top of solidified 20 mL LB agar (1.5% w/v) in a 10 cm Petri dish (100×15 mm, VWR, Cat#89038-968). 100 µL SPβcp phage (∼7.0 × 10^6^ PFUs/mL in LB, or fresh LB alone used as negative control) was spread evenly over the agar surface using an L-shaped spreader. After the surface dried, a 7 mm-diameter filter paper was spotted with 2 µL inhibitor in DMSO and placed on the agar in the center of the Petri dish using sterile forceps. The filter paper was pressed gently to ensure full contact with the agar surface. The plate was incubated at 30 °C for 12-18 hours.

### Measurement of bacterial growth under high-MOI-infection

An overnight culture of +SpbK *B. subtilis* (CMJ82) or –SpbK *B. subtilis* (CU1050) cells (positive control for lysis) was diluted 1:50 into 10 mL fresh LB. The diluted cultures were incubated at 37 °C, 220 rpm for 3 ∼ 4 hours until OD_600nm_ ∼ 0.4. To each well of 96-well plate, 0.2 µL of each inhibitor (100 mM in DMSO) (pure DMSO used as negative control), 100 µL culture and 100 µL SPβcp phage (∼ 1.4 × 10^9^ PFUs/mL) was added in sequence. The plate was incubated at 30 °C, 282 cpm (3 mm orbital shaking) and the OD_600nm_ in each well was recorded every 5 mins using a Synergy H1 plate reader (Biotek).

### Preparation of phage-infected cell lysate for intracellular NAD^+^ level analysis via HPLC-HRMS

Lysates were prepared as described previously^22^, with minor modifications described below. An overnight culture of +SpbK *B. subtilis* (CMJ82) or –SpbK *B. subtilis* (CU1050) (positive control for lysis) was diluted 1:50 into 30 mL fresh LB. The diluted cultures were incubated at 37 °C, 220 rpm for 3 ∼ 4 hours until OD_600nm_ ∼ 0.4. To the 15 mL centrifuge tube containing 4 µL compound 1, 2, or 3 (100 mM in DMSO), 2 mL of the culture was added. Then 2 mL of SPβcp phage (∼ 1.4 × 10^9^ PFUs/mL) was added to the host cells to reach MOI > 10. The infected cell culture was incubated at 30 °C, 220 rpm for 45 mins. After incubation, all samples were immediately centrifuged at 4,200 × g, 4 °C for 5 min. The supernatant was discarded, and the cell pellet was resuspended in 500 µL of 100 mM sodium phosphate buffer (pH = 7) + 4 mg/mL lysozyme. After incubation at room temperature for 10 min, the cells were transferred into 2 mL tubes with Lysing Matrix B (MP Biomedicals #116911050) and lysed using an Omni Bead Ruptor 12 for 2 × 40 s at 6 m/s with a dwell time of 4 min in between. After lysis, all tubes were centrifuged at 14,000 × g, 4 °C for 5 min. Then, 250 µL of each supernatant was transferred to Amicon Ultra-0.5 Centrifugal Filter Units 3 kDa (EMD Millipore #UFC500396) and centrifuged for 20 min at 13,500 × g, 4 °C. The filtrate was collected, concentrated via lyophilization and 10 µL of each was used for LC-MS analysis.

### LC-MS/MS analysis of NAD^+^ in phage-infected cell lysate

Liquid chromatography analysis was performed on an ACQUITY UPLCI-Class PLUS System using a Luna Omega 5-μm Polar C18 100 Å column (250 × 4.6 mm). The mobile phase A was water + 0.1 % (v/v) formic acid, and the mobile phase B was acetonitrile + 0.1 % (v/v) formic acid. The flow rate was kept at 0.7 mL/min and the gradient was as follows: 0% B (0–10 min), increase to 2.5% B (10–15 min), increase to 5% B (15–16 min), hold 5% B (16–26 min), increase to 95% B (26–27 min), hold 95% B (27–37 min), decrease to 0% B (37–38 min), hold 0% B (38–48 min). High-resolution electrospray ionization (HR-ESI) mass spectra with collision-induced dissociation (CID) MS/MS were obtained using a Waters Synapt G2S Quadrupole Time-of-Flight (QTOF). The instrument was operated in negative ionization mode. The MS spectra were obtained on the Time-of-Flight analyzer with a scan range of 300–900 Da and analyzed using MassLynx 4.1 software. The m/z of interest was filtered through the quadrupole, subjected to CID (energy ramp 34–44 V), and analyzed on the Time-of-Flight analyzer with a scan range of 50–750 Da.

### Synergy checkerboard assay

The inhibitor 1 or 2 (100 mM in DMSO) was serially diluted in fresh LB into 4 × final desired concentration. 10 µL of each diluted inhibitor (1 and 2) was separately added into wells of a 384-well plate covering the range of concentrations reported in Fig.4a. An overnight culture of SpbK-expressing *B. subtilis* CU1050 (CMJ82) or no-system CU1050 cells (positive control for lysis) was diluted 1:50 into 10 mL fresh LB. 10 µL of diluted cultures were added to the wells. The plate was pre-incubated at 30 °C for 1 hour, followed with addition of 10 µL SPβcp phage (∼1.4 × 10^6^ PFUs/mL in LB) to each well. The plate was incubated at 30°C in a Biospa8 (Biotek) and the OD_600nm_ in each well was recorded every 1 hour using a Synergy H1 plate reader (Biotek). The inhibition strength of each combination between inhibitor 1 and 2 was calculated described above in section ‘Evaluation of SpbK inhibition in liquid media cultures.’

### Protein production and purification

Both proteins, MBP-SpbK and YonE were expressed and purified as reported by Mishra *et al*.^22^ Specifically, the corresponding plasmids were transformed into *E. coli* BL21(DE3) (New England Biolabs, C2527I) via heat shock. An overnight of transformants was 1:100 diluted into 800 mL LB with ampicillin (100 µg/mL) or kanamycin (50 µg/mL) and incubated at 37 °C, 220 rpm until OD_600nm_ achieved ∼0.8-1.0 (∼4-6 hours). Then the culture was cooled down to 18 °C, and protein expression was induced with 0.6 mM isopropyl-β-D-thiogalactoside (ITPG) overnight at 18 °C.

To purify the MBP-SpbK or YonE, the cell pellet was resuspended in lysis buffer (50 mM HEPES pH 7.5, 30 mM imidazole, 500 mM NaCl) and lysed by sonication. The lysate was centrifuged at 13,000 × g, 8 °C for 20 mins to remove the cell debris. The supernatant was loaded onto a HisTrap™ HP His tag protein purification column (Cytiva, catalog 17524801) pre-equilibrated with wash buffer (50 mM HEPES pH 7.5, 30 mM imidazole, 500 mM NaCl). Then the proteins were eluted with elution buffer (50 mM HEPES pH 7.5, 250 mM imidazole, 500 mM NaCl). The eluted proteins were concentrated with 30 kDa MWCO, Amicon centrifuge filters (Millipore, UFC9010) and loaded onto Superdex 200 pg resin in a HiLoad column, 16/600 (Cytiva, catalog 28989335) pre-equilibrated with buffer (10 mM HEPES pH 7.5, 150 mM NaCl). The eluted protein was concentrated in sequence with 30 kDa MWCO, Amicon centrifuge filters (Millipore, UFC9010) and Amicon Ultra-0.5 Centrifugal Filter Units 10 kDa (Millipore # UFC501096), 4:1 diluted with 50% glycerol and stored at -80 °C. The purity was analyzed by SDS-PAGE.

### HPLC-HRMS-based component analysis of SpbK-mediated NAD^+^ depletion

The reaction was performed in deuterated (∼95%) buffer (50 mM HEPES, 150 mM NaCl in D_2_O, pH = ∼7.5). To a 1.7 mL Eppendorf tube containing 12.5 µL inhibitor (100 mM in DMSO), 200 µL (2.5 µM MBP-SpbK + 37.5 µM YonE in reaction buffer) was added, followed by 12.5 µL reaction buffer and 25 µL NAD^+^ (5 mM in reaction buffer). After complete mixing via vortex, the tube was incubated at 30 °C.

To prepare the samples for HPLC-HRMS analysis, 50 µL of the above reaction mixture was transferred into a 1.7 mL Eppendorf tube. After heating to 95 °C for 3 min to stop the reaction and subsequent centrifugation at 20,000 × g at 4°C for 5 min to remove the precipitated protein, 10 µL of the supernatant was analyzed by HPLC-MS.

The HPLC-HRMS settings are described above in the section ‘LC-MS/MS analysis of NAD^+^ in phage-infected cell lysate’. The gradient was changed as follows: 0% B (0–10 min), increase to 2.5% B (10–15 min), increase to 5% B (15–16 min), hold 5% B (16-26 min), increase to 95% B (26–36 min), hold 95% B (36–46 min), decrease to 0% B (46–47 min), hold 0% B (47–57 min). Collision-induced dissociation (CID) MS/MS was set up to collected based on specified m/z of compound-ADPR conjugates.

### NMR-based evaluation of generation of inhibitor-ADPR conjugates

The sample prepared in the above section ‘HPLC-HRMS-based component analysis of SpbK-mediated NAD^+^ depletion’ was transferred to a 3 mm Bruker NMR tube. All ^1^H NMR spectra were acquired with a Bruker 800 MHz Avance Neo spectrometer equipped with 5 mm Triple Resonance TCI CryoProbe with cooled preamps. To further suppress resonance from H_2_O, a water suppression pulse program (P3919GP) was implemented.

### Computational modeling

To model and compare possible bound structures of SpbK inhibitors, Boltz-2^66^ was employed to co-fold SpbK with each of the selected inhibitors and native ligands, respectively. An AlphaFold 3 model of hexametric SpbK TIR domains^22^ was used as the reference. Its corresponding sequence (Glu-111 to Lys-266) in hexametric form and the SMILES string of each ligand (monomeric) were used as molecular input in Boltz-2 YAML files. A series of distance constraints were applied in Boltz-2 to maintain the hexametric protein structure, which ensures integrity of the composite active site at TIR-TIR interface, and proximity of the inhibitor ligand towards the active site. These include 15 distance constraints for each pair of the six Glu-192 residues from the hexamer, with the maximum distance value one angstrom larger than the pairwise CA distance measured in the AlphaFold 3 model to allow flexibility, as well as another distance constraint between each of the ligand and Glu-192 in chain C of the hexamer, with the maximum distance value at 6 angstroms. Co-folding calculations were performed on a high-performance computing cluster at Monash University with NVIDIA L40S GPU. Most command-like interface parameters were set to default values, except for the recycling step (set to 5) and the sampling step (set to 200). Due to the stochastic nature of Boltz-2 sampling, co-folding of each protein-ligand pair was run in triplicates and one representative prediction (with the exchanged base on the same side of the adjacent ribose ring as in the AlphaFold 3 model and the ribose interacting with Glu-192) was selected for analysis.

### Evaluation of SpbK inhibition and SPβ phage production in SPβ -lysogenized bacteria

An overnight culture of +ICE*Bs1 B. subtilis* (168) or –ICE*Bs1 B. subtilis* (MB8_B7) cells (positive control for lysis) was diluted 1:100 into 20 mL fresh LB. The diluted cultures were transferred into wells of a flat-bottom 96-well plate. The plate was incubated at 37 °C, 282 cpm (3 mm orbital shaking) and the OD_600nm_ in each well was recorded every 5 mins using a Synergy H1 plate reader (Biotek). When the OD_600nm_ reached ∼ 0.4 (mid-exponential phase) (3 ∼ 4 hours), one of the following solutions was added to total final volume of 200 µL. (i) Milli-Q water (untreated control); (ii) mitomycin C (MMC), 0.4 µg/mL final concentration; (iii) compound 8 (Ambeed, Cat#A344521), 100 µM final concentration; or (iv) compound 8 (100 µM) + MMC (0.4 µg/mL). All stock solutions were prepared in Milli-Q water. Each treatment was performed with 7 biological replicates for growth curve analysis; a subset of 3 biological replicates (each with 2 technical replicates) was used for subsequent SPβ phage production measurement.

The above samples for measuring SPβ phage production were collected 2.5 h after treatment. The cultures were transferred to microcentrifuge tubes and centrifuged at 8,000 × g for 5 min at room temperature to pellet cell debris. The clarified supernatant was passed through a 0.22 µm filter to remove residual cells and large particles. Filtered supernatants were immediately subjected to serial ten-fold dilutions in saline solution (0.9 % NaCl), and 2.5 µL of each dilution was spotted in triplicate onto lawns on indicator strain *B. subtilis* Δ6.

For lawn preparation, 300 µL of the exponentially growing *B. subtilis* Δ6 culture (OD_600nm_ ∼ 0.4) was added to 12 mL of LB soft agar (0.3% agar, w/v) equilibrated at 60 °C, mixed gently, and immediately poured onto square polystyrene Petri dishes (120 × 120 × 17 mm, Greiner Austria, 688102). Plates were allowed to cool and solidify at room temperature for at least 20 min before spotting. Filtered supernatants and serial dilutions were spotted as 2.5 µL drops and allowed to absorb into the overlay for 10–15 min before the plates were inverted and incubated overnight at 28 °C. Plaques were counted the following day and phage titers were expressed as plaque-forming units per milliliter (PFU/mL), calculated as:

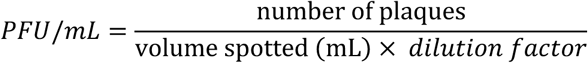

## Resource availability Lead contact

Requests for further information and resources should be directed to and will be fulfilled by the lead contact, Joseph P. Gerdt.

## Data availability

Any requests for data should be addressed to the corresponding author.

## Materials & Correspondence

Correspondence and requests for materials should be addressed to Joseph P. Gerdt.

## Acknowledgments

We thank the National Cancer Institute Development Therapeutics Program (NCI/DTP) for providing compounds present in this manuscript, specifically, the Diversity Library Set VII, compounds 1 (NSC 70717), 2 (NSC 63001), 3 (NSC 204920), 1-5 (NSC 102086) and 1-6 (NSC 131986). We thank Alan Grossman (Massachusetts Institute of Technology) for providing bacteria and phages. We thank Xinfeng (Frank) Gao and Kathrine Ann Edmonds (NMR facility, Indiana University Bloomington) for support with NMR experiments. We thank Jonathan Cyboski Trinidad (MS facility, Indiana University Bloomington) for support with HPLC-HRMS experiments. This work was also supported by Monash eResearch capabilities, including High Performance Computing (HPC). The research was funded by a National Science Foundation CAREER award (IOS-2143636) to J.P.G. and a Camille Dreyfus Teacher-Scholar Award (TC-24-028) to J.P.G. A.D and J.J.L were supported by European Union (ERC, PHAGECONTROL, 101041421). Y.S. was supported by an Australian Research Council Discovery Early Career Research Award (DE250101258). The 800 MHz NMR spectrometer used in this research was purchased in part with support from the Lilly Endowment and NIH award S10 OD032431-01A1.

## Author contributions

Conceptualization, Y.C., J.P.G.; methodology, Y.C., J.J.L., Y.S., A.D., J.P.G.; analysis, Y.C., J.J.L., Y.S., A.D., J.P.G.; investigation, Y.C., J.J.L., Y.S., B.P.M., A.D., J.P.G.; writing – original draft, Y.C.; writing – review & editing, Y.C., J.J.L., Y.S., T.V., A.D., J.P.G.; visualization, Y.C., J.J.L., Y.S., A.D., J.P.G.; supervision, Y.S. T.V., A.D., J.P.G.; funding acquisition, Y.S. T.V., A.D., J.P.G.

---

**Figure S1.**
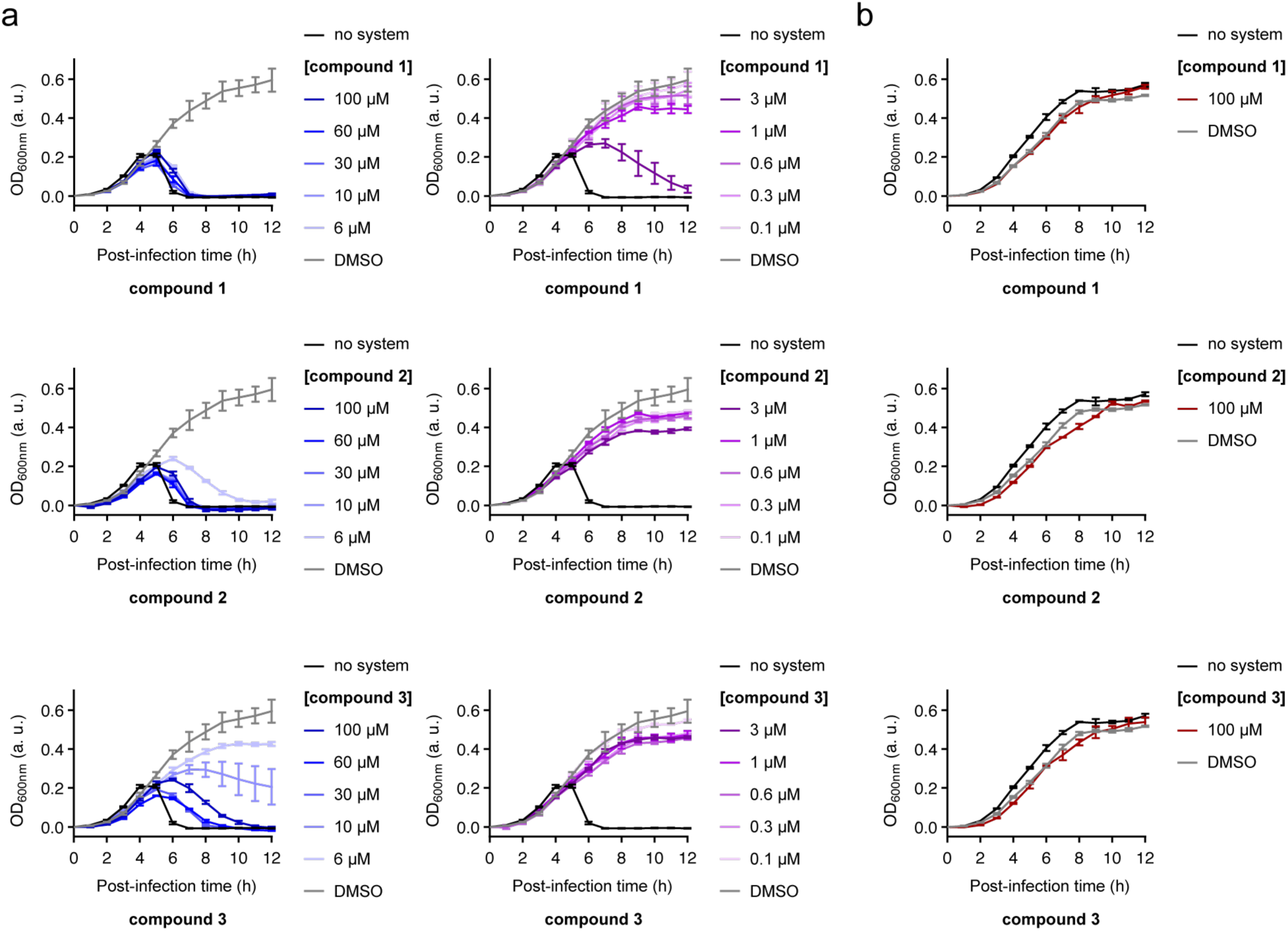
Compounds 1–3 inhibit SpbK. (a) Lysis curves of +SpbK strain treated with SPβcp phage (MOI = 0.3) and different concentrations of **compound 1 or 2 or 3** compared to a control (–SpbK strain, black). (b) Lysis curves of +SpbK strain treated with tested maximum concentration (100 µM) of **compound 1 or2 or 3** compared to a control (–SpbK strain, black). In both panels a & b, data are represented as the mean ± SEM of n = 3 independent biological replicates.

**Figure S2.**
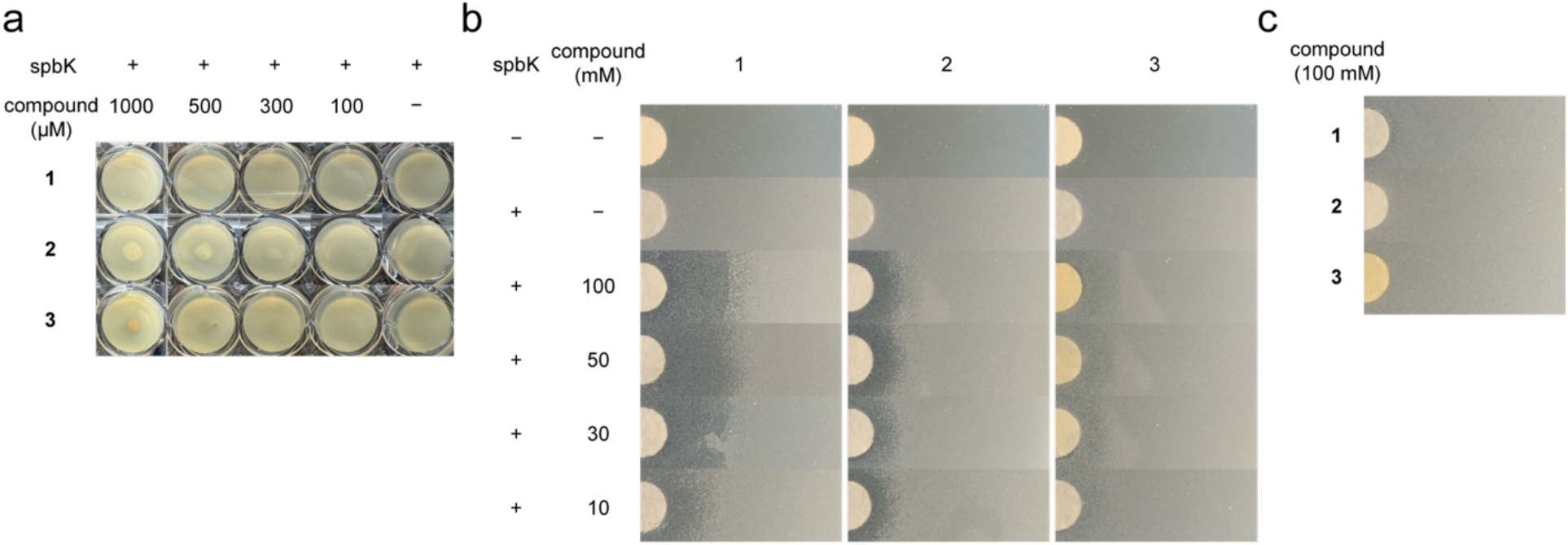
Inhibitors selectively promote phage-induced lysis. (a) 10 µL Lysogeny-Broth medium was spotted onto +SpbK *B. subtilis* (CMJ82) with different concentrations of **compounds 1–3**, compared to a control (DMSO, last column). No growth defects were observed in the absence of phage. (b) 2 µL compound (serial concentrations in DMSO) spotted onto 7 mm-diameter filter paper lying on +SpbK *B. subtilis* (CMJ82) under SPβcp infection compared to a control (+SpbK *B. subtilis* [CMJ82]). (c) 2 µL compound (100 mM in DMSO) spotted onto a 7 mm-diameter filter paper lying on +SpbK *B. subtilis* (CMJ82). No growth defects were observed in the absence of phage.

**Figure S3.**
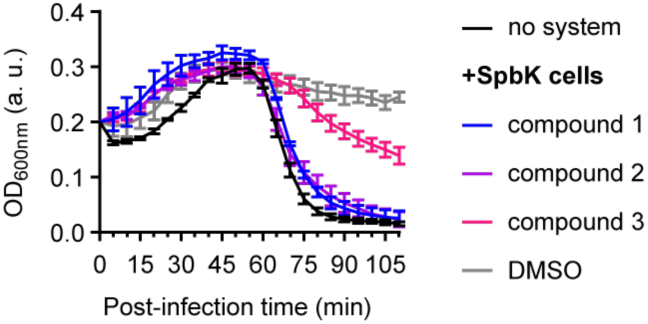
Inhibitors promote High-MOI-phage-induced lysis in liquid media culture. Lysis curves of +SpbK *B. subtilis* (CMJ82) (–SpbK *B. subtilis* [CU1050] used as control, black) treated with SPβcp phage (MOI ∼ 10) and 100 µM inhibitor 1, 2, and 3 (DMSO used as control, grey).

**Figure S4.**
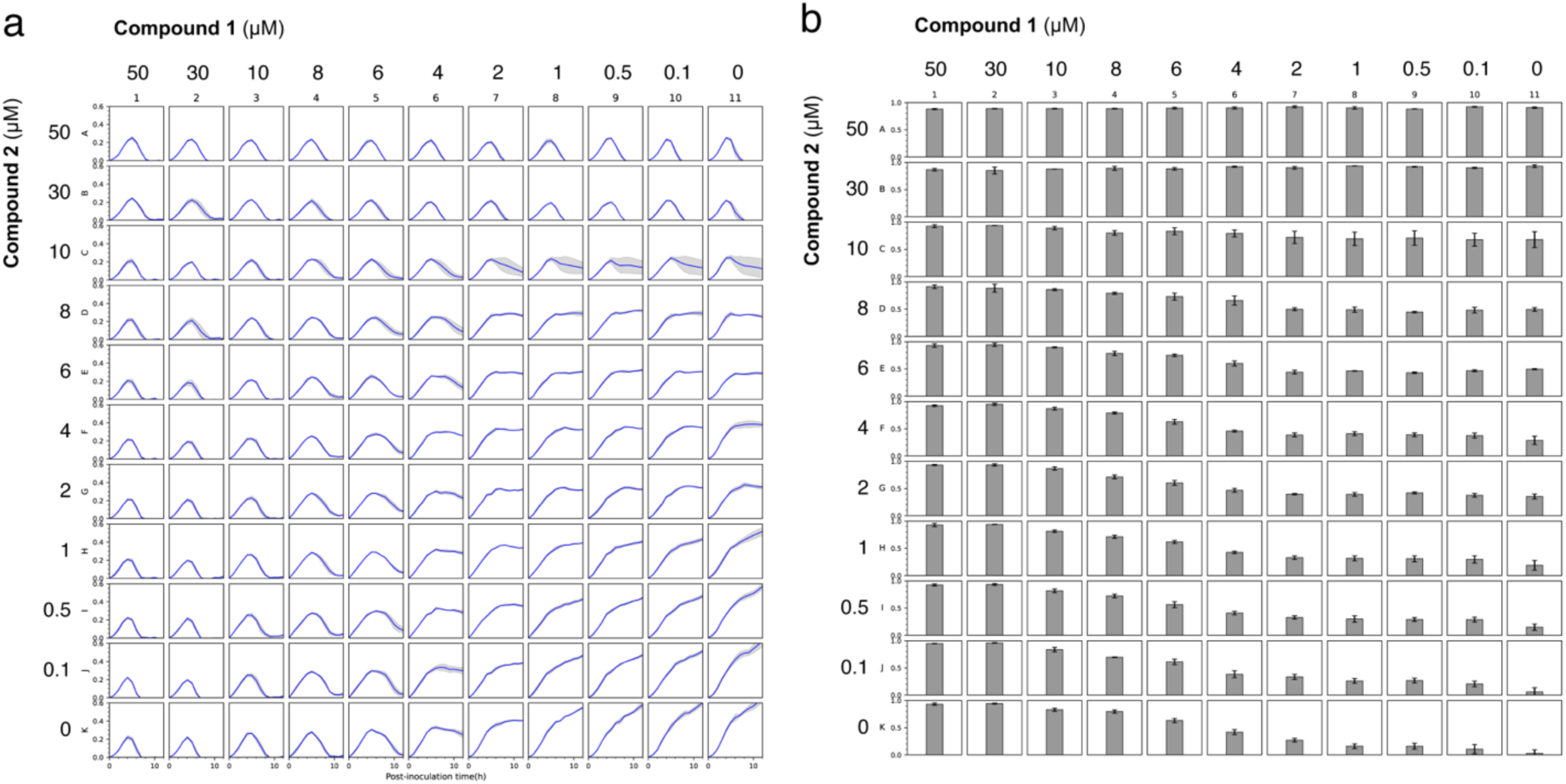
Compounds 1 and 2 exhibit additive inhibition. (a) Lysis curves of +SpbK strain treated with SPβcp phage (MOI = 0.3) and different concentration combinations of compound 1 and 2 in the synergy checkerboard assay. Data are normalized and represented as the mean (blue curve) ± SEM (grey area) of n = 3 independent biological replicates. (b) SpbK inhibition strength of each concentration combination of compound 1 and 2 presented as the mean (grey column) ± SEM (black error bar) of n = 3 independent biological replicates.

**Figure S5.**
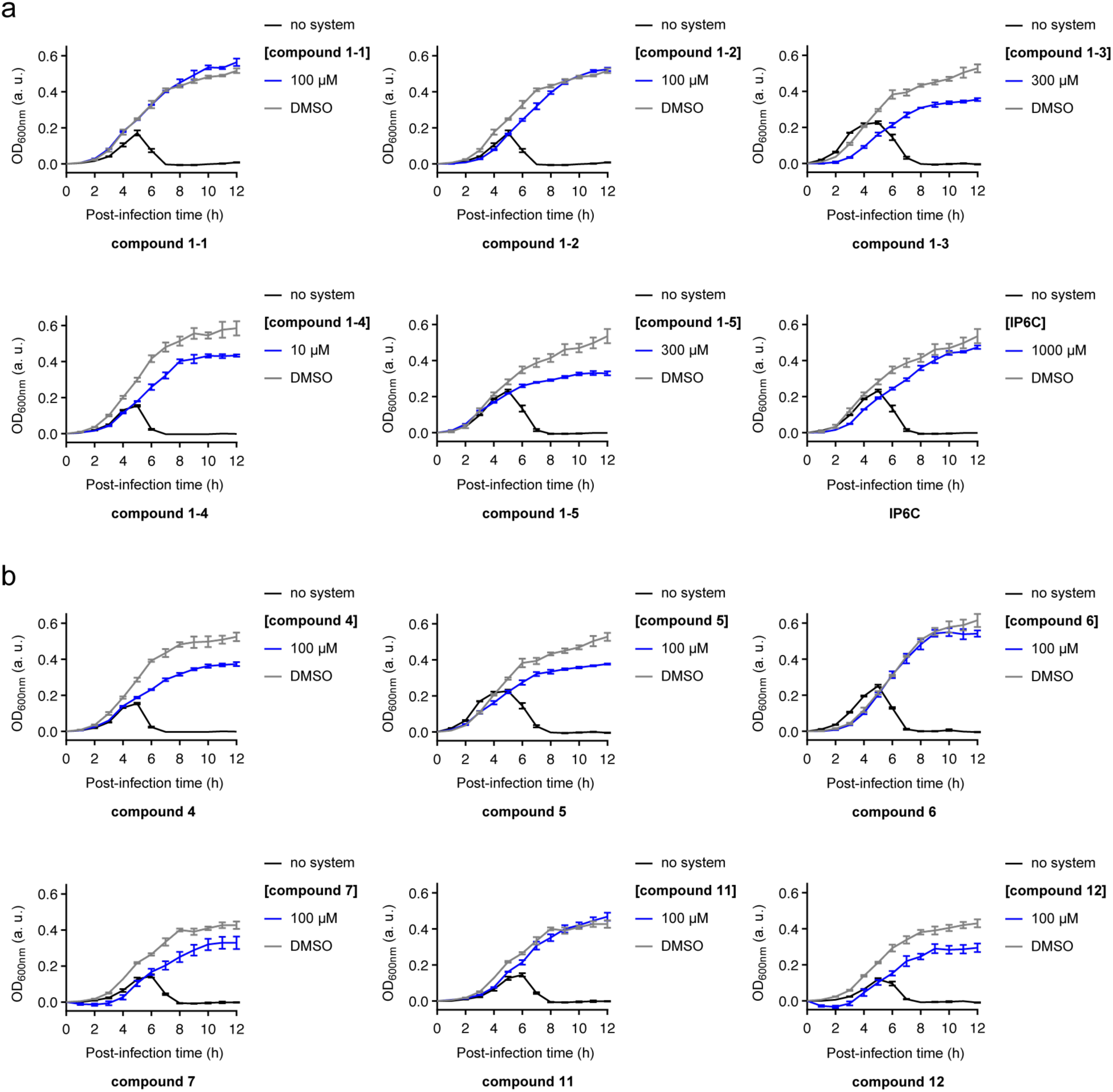
Inactive compounds. (a) Lysis curves of +SpbK strain treated with SPβcp phage (MOI = 0.3) and the maximum tested non-cytotoxic concentrations of **compounds 1-1** to **1-5** and **IP6C** compared to a control (–SpbK strain, black). Each compound was tested up to 1 mM concentration. (b) Lysis curves of +SpbK strain treated with SPβcp phage (MOI = 0.3) and the maximum tested concentration (100 µM) of **compounds 4 - 7, 11** and **12** compared to a control (–SpbK strain, black). In both panels a & b, data are represented as the mean ± SEM of n = 3 independent biological replicates.

**Figure S6.**
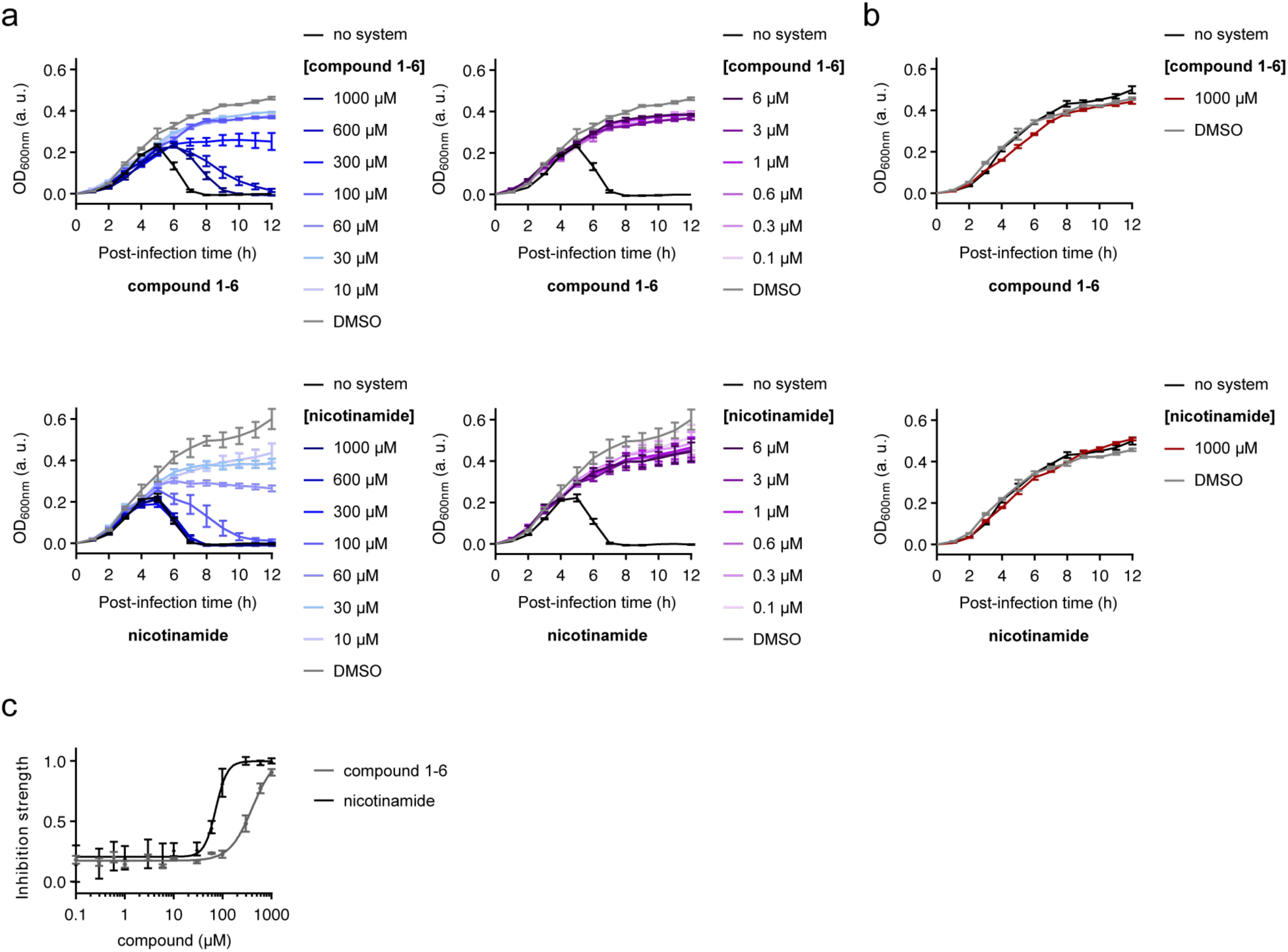
Compound 1-6 and nicotinamide inhibit SpbK. (a) Lysis curves of +SpbK strain treated with SPβcp phage (MOI = 0.3) and different concentrations of **compound 1-6** or **nicotinamide** compared to a control (–SpbK strain, black). (b) Lysis curves of +SpbK strain treated with the maximum tested concentration (1 mM) of **compound 1-6** or **nicotinamide** compared to a control (–SpbK strain, black). (c) Dose-response curves of **compound 1-6** and **nicotinamide** measured with +SpbK strain treated with SPβcp phage (MOI = 0.3). In all panels, data are represented as the mean ± SEM of n = 3 independent biological replicates.

**Figure S7.**
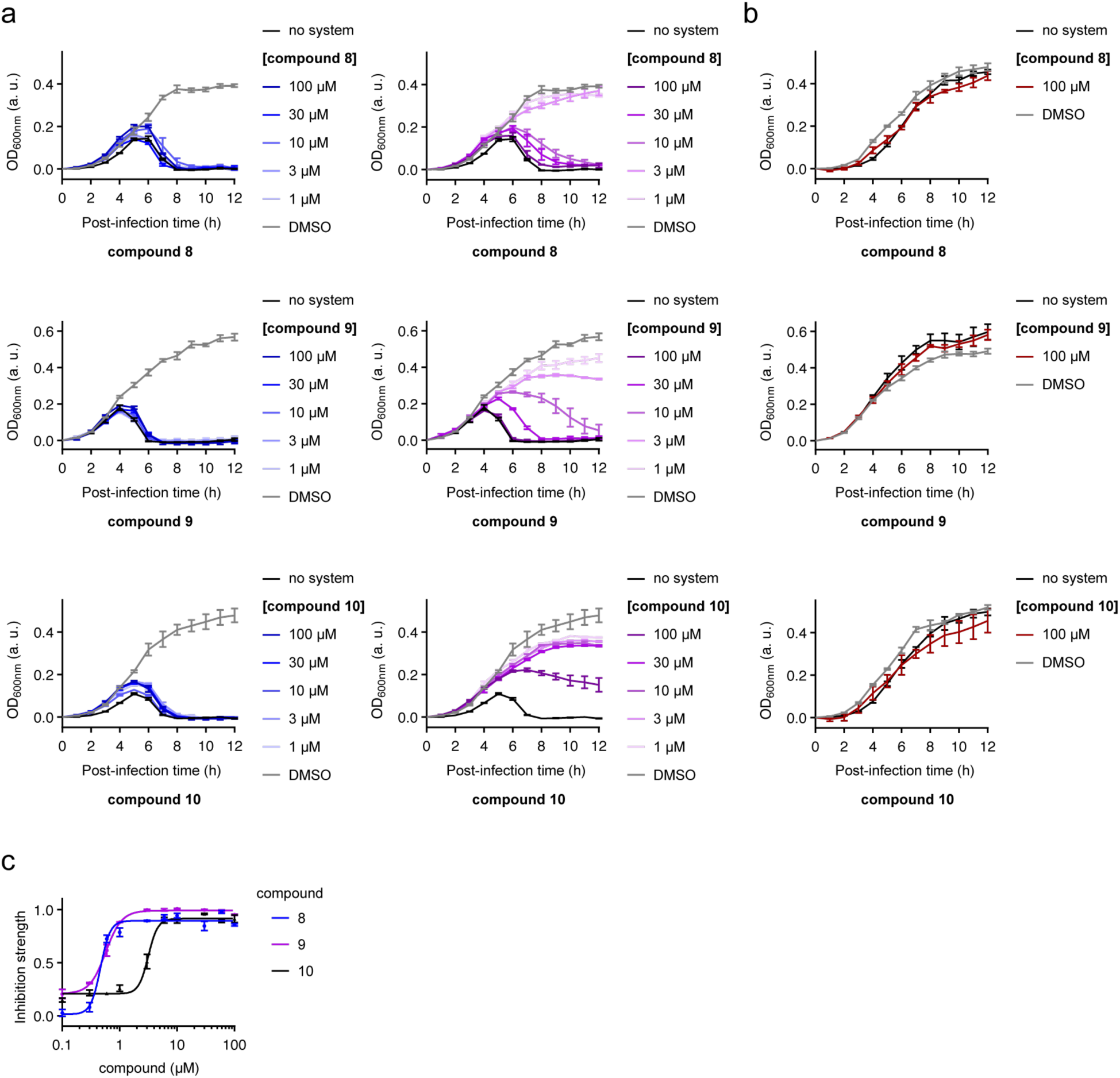
Compounds 8–10 inhibit SpbK. (a) Lysis curves of +SpbK strain treated with SPβcp phage (MOI = 0.3) and different concentrations of **compound 8** or **9** or **10** compared to a control (–SpbK strain, black). (b) Lysis curves of +SpbK strain treated with the maximum tested concentration (100 µM) of **compound 8** or **9** or **10** compared to a control (–SpbK strain, black). (c) Dose-response curves of **compound 8**, **9** and **10** measured with +SpbK strain treated with SPβcp phage (MOI = 0.3). In all panels, data are represented as the mean ± SEM of n = 3 independent biological replicates.

**Figure S8.**
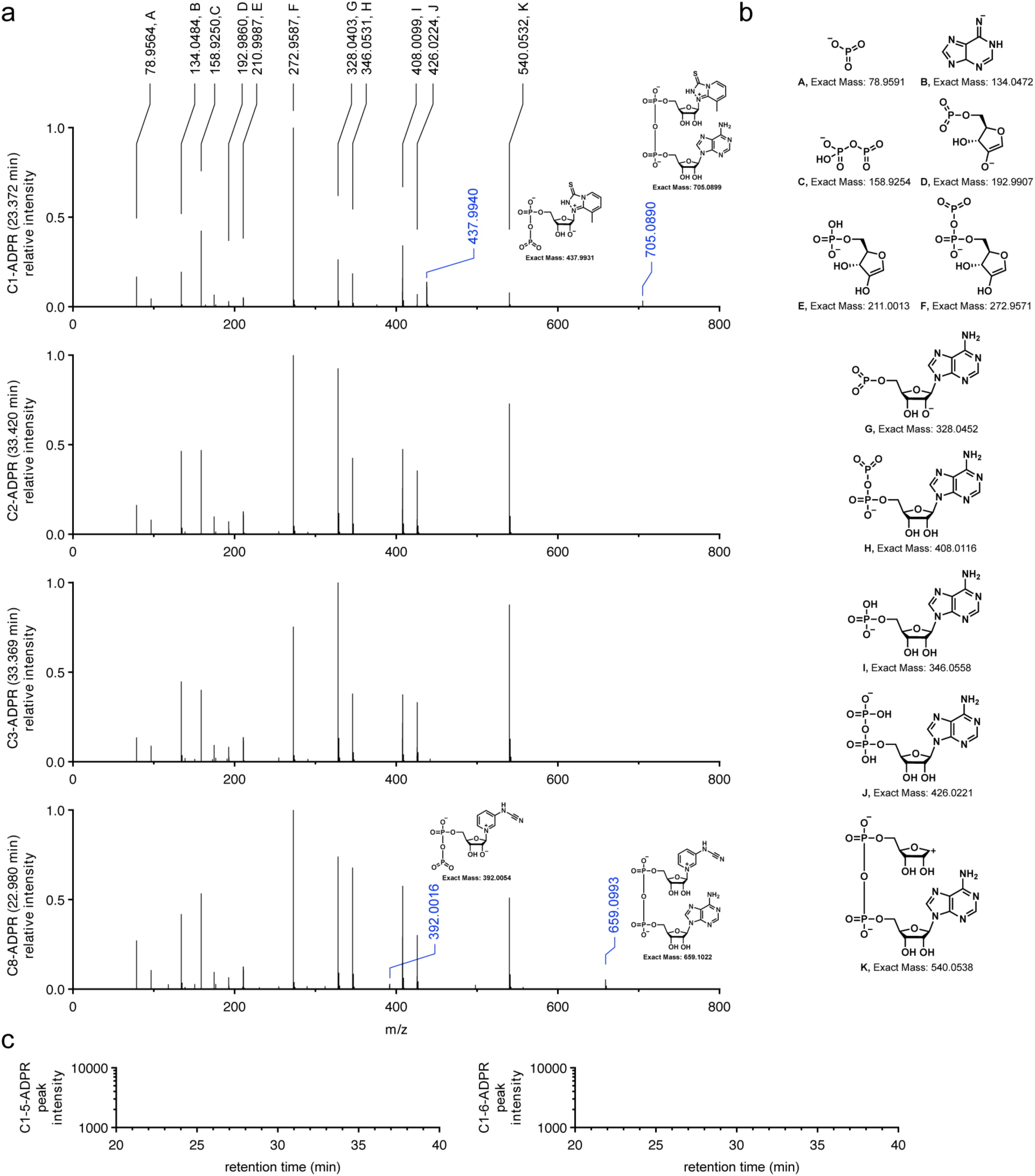
HPLC-HRMS-based detection of generation of compound-ADPR conjugates. (a) MS/MS spectra of the compound-ADPR conjugates reported in Fig. 3c. The shared characteristic m/z’s generated from ADPR moiety of conjugate are labeled in black. Unique characteristic fragment m/z’s are labeled in blue with their corresponding predicted chemical structures aside. (b) Predicted chemical structures of characteristic ADPR fragmentation patterns indicated in Fig. S8a. (c) Extracted ion chromatogram (EIC) for m/z 777.0955-777.1265 (matching C1-5-ADPR [M-H]^-^) and m/z 748.1423-748.1723 (matching C1-6-ADPR [M–H]^-^). No peak with higher intensity than 1000 was detected.

**Figure S9.**
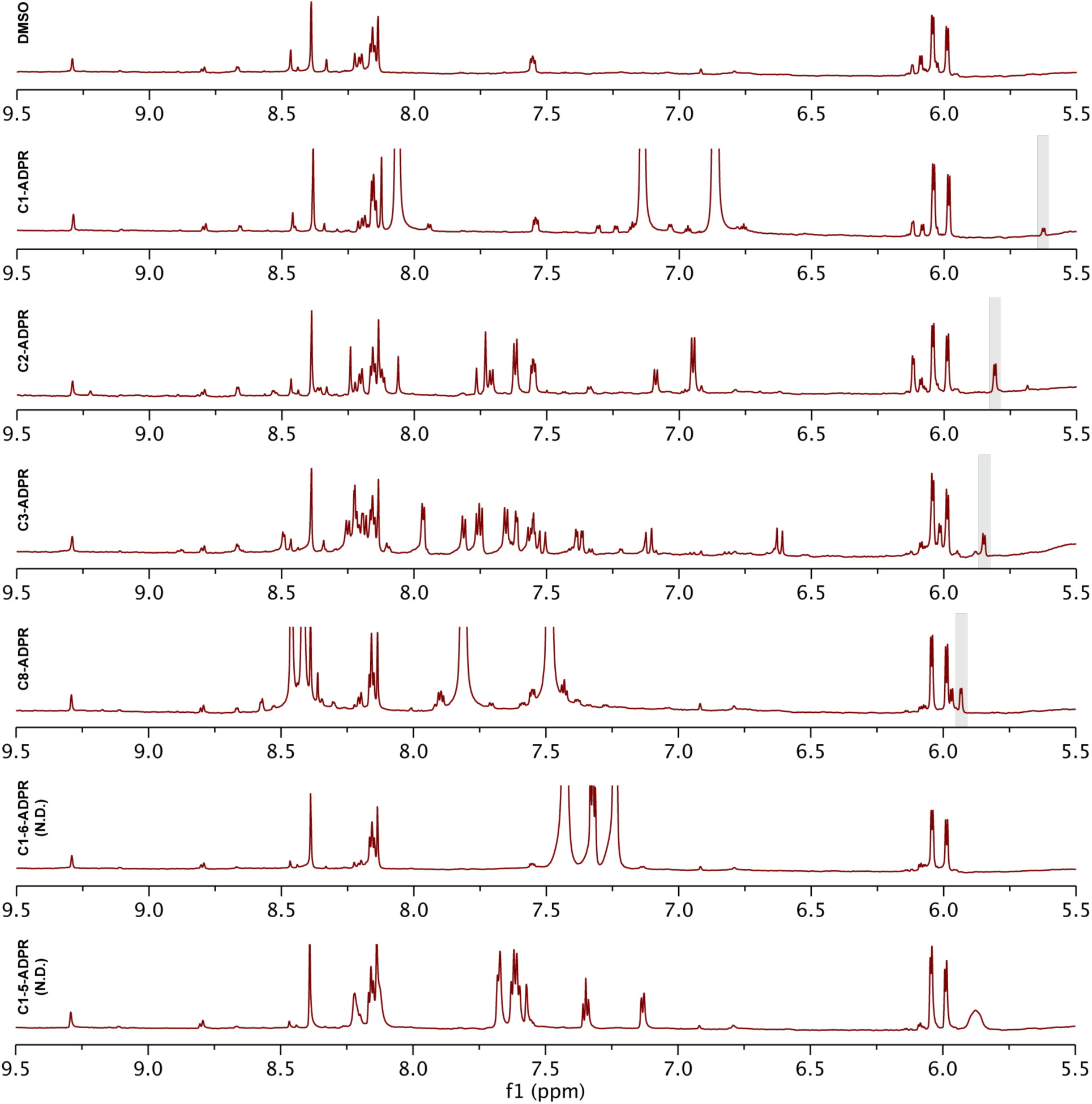
**^1^H-NMR-based detection of generation of compound-ADPR conjugators.**^1^H-NMR spectra (5.5-9.5 ppm) of each reaction mixture at 12 hours in Fig. 3b. The peak of hydrogen connected to ribose anomeric carbon in each compound-ADPR conjugator was indicated by grey shadow. N.D. means corresponding ADPR conjugate was not detected.

**Table S1.** Predicted binding affinities

| Ligand | IC<sub>50</sub> (μM) | Log<sub>10</sub>[IC<sub>50</sub>(μM)] |
| --- | --- | --- |
| NAD<sup>+</sup> | 4.21 | 0.62 |
| N1-cADPR | 15.5 | 1.19 |
| C1-ADPR | 6.98 | 0.84 |
| C2-ADPR | 2.28 | 0.36 |
| C3-ADPR | 0.46 | -0.34 |
| C8-ADPR | 3.32 | 0.52 |
| C1-5 | 30.0 | 1.48 |
| C1-6 | 19.5 | 1.29 |

Predicted affinities of ligands for SpbK TIR domain (residues: 113-266) hexamer (PDB:9PHB) by co-folding using Boltz-2.

**Table S2.** Strain information of *Bacillus subtilis* used in the genetic analysis

| *B. subtilis* strain | Accession | ICEBs1 | *spbK* | SPβ-like | *yonE* |
| --- | --- | --- | --- | --- | --- |
| *Bacillus subtilis* SRCM100761 | CP021889.1 | yes | - | - | - |
| *Bacillus subtilis* SRCM103581 | CP035403.1 | yes | - | - | - |
| *Bacillus subtilis* PN176 (HK176) | CP103456.1 | yes | - | - | - |
| *Bacillus subtilis* BEST195 | AP011541.2 | yes | - | - | - |
| *Bacillus subtilis* CGMCC 2108 | CP014471.1 | yes | - | - | - |
| *Bacillus subtilis* BS38 | CP017314.1 | yes | - | - | - |
| *Bacillus subtilis* 29R7-12 | CP017763.1 | yes | - | - | - |
| *Bacillus subtilis* KH2 | CP018184.1 | yes | - | - | - |
| *Bacillus subtilis* ATCC 21228 | CP020023.1 | yes | - | - | - |
| *Bacillus subtilis* Bs-115 | CP020722.1 | yes | - | - | - |
| *Bacillus subtilis* SRCM101444 | CP021498.1 | yes | - | - | - |
| *Bacillus subtilis* SRCM100757 | CP021499.1 | yes | - | - | - |
| *Bacillus subtilis* MH-1 | CP032853.1 | yes | - | - | - |
| *Bacillus subtilis* N1-1 | CP032861.1 | yes | - | - | - |
| *Bacillus subtilis* N2-2 | CP032863.1 | yes | - | - | - |
| *Bacillus subtilis* N3-1 | CP032865.1 | yes | - | - | - |
| *Bacillus subtilis* N4-2 | CP032867.1 | yes | - | - | - |
| *Bacillus subtilis* SRCM103576 | CP035402.1 | yes | - | - | - |
| *Bacillus subtilis* SRCM103612 | CP035406.1 | yes | - | - | - |
| *Bacillus subtilis* SRCM103622 | CP035411.1 | yes | - | - | - |
| *Bacillus subtilis* CMIN-4 | CP063151.1 | yes | - | - | - |
| *Bacillus subtilis* JNFE1126 | CP070485.1 | yes | - | - | - |
| *Bacillus subtilis* BN0 | CP070844.1 | yes | - | - | - |
| *Bacillus subtilis* JNFE0126 | CP071276.1 | yes | - | - | - |
| *Bacillus subtilis* SCP010-1 | CP086007.1 | yes | - | - | - |
| *Bacillus subtilis* SPC-17-3 | CP098036.1 | yes | - | - | - |
| *Bacillus subtilis* NB205 | CP098491.1 | yes | - | - | - |
| *Bacillus subtilis* BGSC 27E1 | CP101932.1 | yes | - | - | - |
| *Bacillus subtilis* Bacillus subtilis 'natto' | CP101933.1 | yes | - | - | - |
| *Bacillus subtilis* SRCM116301 | CP106671.1 | yes | - | - | - |
| *Bacillus subtilis* SRCM116268 | CP106673.1 | yes | - | - | - |
| *Bacillus subtilis* 7PJ-16 | CP023409.1 | yes | - | yes | - |
| *Bacillus subtilis* GL-4 | CP104097.1 | yes | - | yes | - |
| *Bacillus subtilis* PS-168 | SAMN36922892 | yes | yes | - | - |
| *Bacillus subtilis* PS-31 | SAMN08637086 | yes | yes | - | - |
| *Bacillus subtilis* PS-68 | SAMN08637090 | yes | yes | - | - |
| *Bacillus subtilis* PS-96 | SAMN36922886 | yes | yes | - | - |
| *Bacillus subtilis* 168 | NC_000964.3 | yes | yes | yes | yes |
| *Bacillus subtilis* NBRC 13719 | AP019714.1 | yes | yes | yes | yes |
| *Bacillus subtilis* 6051-HGW | CP003329.1 | yes | yes | yes | yes |
| *Bacillus subtilis* KCTC 1028 | CP011115.1 | yes | yes | yes | yes |
| *Bacillus subtilis* KCTC 3135 | CP015375.1 | yes | yes | yes | yes |
| *Bacillus subtilis* FJAT-4 | CP082924.1 | yes | yes | yes | yes |
| *Bacillus subtilis* Bsi | CP089592.1 | yes | yes | yes | yes |
| *Bacillus subtilis* YZSR384 | CP102769.1 | yes | yes | yes | yes |
| *Bacillus subtilis* 137 | CP103066.1 | yes | yes | yes | yes |
| *Bacillus subtilis* P8_B3 | CP045812.1 | yes | yes | yes | yes |
| *Bacillus subtilis* P8_B1 | CP045922.1 | yes | yes | yes | yes |
| *Bacillus subtilis* O4 | CP112873.1 | yes | yes | yes | yes |
| *Bacillus subtilis* N1282-4at | CP064093.1 | yes | yes | yes | yes |
| *Bacillus subtilis* N1108-5at | CP064094.1 | yes | yes | yes | yes |
| *Bacillus subtilis* N1303-2Ay | CP064095.1 | yes | yes | yes | yes |
| *Bacillus subtilis* N1142-3at | CP064096.1 | yes | yes | yes | yes |
| *Bacillus subtilis* SZMC 6179J | CP015004.1 | - | - | yes | yes |
| *Bacillus subtilis* 2014-3557 | CP045672.1 | - | - | yes | yes |
| *Bacillus subtilis* MB8_B7 | CP045821.1 | - | - | yes | yes |
| *Bacillus subtilis* PS-14 | SAMN36922878 | - | - | yes | yes |
| *Bacillus subtilis* PS-233 | SAMN36922894 | - | - | yes | yes |
| *Bacillus subtilis* D12-5 | CP014858.1 | - | - | yes | yes |
| *Bacillus subtilis* HJ0-6 | CP016894.1 | - | - | yes | yes |
| *Bacillus subtilis* NCD-2 | CP023755.1 | - | - | yes | yes |
| *Bacillus subtilis* H1 | CP026662.1 | - | - | yes | yes |
| *Bacillus subtilis* SRCM102754 | CP028202.1 | - | - | yes | yes |
| *Bacillus subtilis* SRCM102748 | CP028212.1 | - | - | yes | yes |
| *Bacillus subtilis* SRCM102749 | CP028213.1 | - | - | yes | yes |
| *Bacillus subtilis* HMNig-2 | CP031784.1 | - | - | yes | yes |
| *Bacillus subtilis* MZK05 | CP032315.1 | - | - | yes | yes |
| *Bacillus subtilis* SSJ-1 | CP032860.1 | - | - | yes | yes |
| *Bacillus subtilis* SRCM103517 | CP035226.1 | - | - | yes | yes |
| *Bacillus subtilis* SRCM103612 | CP035406.1 | - | - | yes | yes |
| *Bacillus subtilis* JAAA | CP045425.1 | - | - | yes | yes |
| *Bacillus subtilis* MB8_B1 | CP045823.1 | - | - | yes | yes |
| *Bacillus subtilis* MB8_B10 | CP045824.1 | - | - | yes | yes |
| *Bacillus subtilis* R31 | CP046591.1 | - | - | yes | yes |
| *Bacillus subtilis* pb2441 | CP068982.1 | - | - | yes | yes |
| *Bacillus subtilis* NIB353 | CP089148.1 | - | - | yes | yes |
| *Bacillus subtilis* ZW | CP092369.1 | - | - | yes | yes |
| *Bacillus subtilis* YB-15 | CP092631.1 | - | - | yes | yes |
| *Bacillus subtilis* N4 | CP095736.1 | - | - | yes | yes |
| *Bacillus subtilis* SRCM117108 | CP103350.1 | - | - | yes | yes |
| *Bacillus subtilis* 4ZT | CP104878.1 | - | - | yes | yes |
| *Bacillus subtilis* MG-1 | CP110634.1 | - | - | yes | yes |
| *Bacillus subtilis* W7 | CP115738.1 | - | - | yes | yes |
| *Bacillus subtilis* 21855 | CP118167.1 | - | - | yes | yes |
| *Bacillus subtilis* PS-11 | SAMN08637079 | - | - | yes | yes |
| *Bacillus subtilis* PS-160 | SAMN08637092 | - | - | yes | yes |
| *Bacillus subtilis* PS-194 | SAMN36922893 | - | - | yes | yes |
| *Bacillus subtilis* PS-261 | SAMN08637099 | - | - | yes | yes |
| *Bacillus subtilis* PS-54 | SAMN36922881 | - | - | yes | yes |
| *Bacillus subtilis* PS-55 | SAMN36922882 | - | - | yes | yes |
| *Bacillus subtilis* SRCM124333 | CP116012.1 | - | - | yes | - |
| *Bacillus subtilis* FX-1 | CP061870.1 | - | - | yes | - |
| *Bacillus subtilis* MB9_B1 | CP045820.1 | - | - | yes | - |
| *Bacillus subtilis* SRCM103881 | CP035165.1 | - | - | yes | - |
| *Bacillus subtilis* SRCM102756 | CP028218.1 | - | - | yes | - |
| *Bacillus subtilis* BS155 | CP029052.1 | - | - | yes | - |
| *Bacillus subtilis* BS16045 | CP017112.1 | - | - | yes | - |

This table lists all strains included in the genetic analysis, along with their accession numbers (GenBank or SAMN, depending on the availability of complete genome sequences). The other columns indicate the presence of the *ICEBsI* element, the *spbK* gene within *ICEBsI*, the SPβ -like prophage, and the *yonE* gene associated with the phage. Only strains containing either *ICEBsI* and/or an SPβ -like prophage were included in the analysis. The presence of these mobile genetic elements (MGEs) was inferred from previous analyses performed by Stefanič *et al.*<sup>1</sup> and from clustering data available on Zenodo<sup>2</sup> associated with that manuscript. The presence of specific genes (*spbK* and *yonE*) was subsequently verified by manual BLAST analysis.

**Table S3.** Bacteria and bacteriophage used in this study

| Strains | Description | Source |
| --- | --- | --- |
| *B. subtilis* CMJ82 | CU1050 (ICEBs1<sup>0</sup>) (SPβ<sup>0</sup>) *amyE:: {spbK cat}* | Johnson *et al.*<sup>3</sup> |
| *B. subtilis* CU1050 | ICEBs1<sup>0</sup>SPβ<sup>0</sup>*metA thrC leu codY sup-3 (trnS-lys)*<sup>4</sup> (CMJ28) | Johnson *et al.*<sup>3</sup> |
| *B. subtilis* 168 | ICEBs1<sup>1</sup>SPβ<sup>1</sup>, accession AL009126/NC_000964.3 | Kunst *et al.*<sup>5</sup> |
| *B. subtilis* MB8_B7 | ICEBs1<sup>0</sup>SPβ<sup>1</sup>, accession NZ_CP045821.1 | Heiko *et al.*<sup>6</sup> |
| *B. subtilis* Δ6 | SPβ indicator strain, accession NZ_CP015975 | Westers *et al.*<sup>7</sup> |
| SPβcp | spontaneous clear plaque mutant of SPβ | Loyo *et al.*<sup>8</sup> |

**Table S4.** Recombinant protein used in this study.

| Recombinant protein | Residue range | Vector | Source |
| --- | --- | --- | --- |
| MBP-SpbK | SpbK: 1-266 | pET-28(b), N-terminal His<sub>6</sub>-tag, TEV protease cleavage site between MBP and SpbK | Mishra *et al.*<sup>9</sup> |
| YonE | YonE (45-506) | pMCSG7, N-terminal His<sub>6</sub>-tag followed by TEV protease cleavage site | Mishra *et al.*<sup>9</sup> |

**Table S5.** Chemicals and commercial assays used in this study

| Chemicals | Source | Identifier |
| --- | --- | --- |
| LB broth | VWR | Cat#90003-350 |
| LB broth | Condalab | Cat#1231 |
| Dehydrated Agar | Fisher | Cat#DF0140-07-4 |
| Kanamycin | Sigma Aldrich | Cat#K1377 |
| Ampicillin | Sigma Aldrich | Cat#A9518 |
| HEPES | Sigma Aldrich | Cat#83264 |
| Imidazole | Sigma Aldrich | Cat#I2399 |
| NaCl | VWR | Cat#BDH9286 |
| DMSO | Sigma Aldrich | Cat#D8418 |
| IPTG | Gold Biotechnology | Cat#I2481C50 |
| Formic acid (HPLC) | Thermo Fisher scientific | Cat#T85178 |
| K<sub>2</sub>HPO<sub>4</sub> | Sigma Aldrich | Cat#S5136 |
| KH<sub>2</sub>PO<sub>4</sub> | Sigma Aldrich | Cat#S3139 |
| Lysozyme | Research Products International | Cat#L38100 |
| NAD<sup>+</sup> | Sigma Aldrich | Cat#N1511 |
| D<sub>2</sub>O | Sigma Aldrich | Cat#151882 |
| d<sub>6</sub>-DMSO | Sigma Aldrich | Cat#151874 |
| Compound 1 | Requested from NCI | NSC70717 |
| Compound 1-1 | Combi-Blocks | Cat#QM-4957 |
| Compound 1-2 | Combi-Blocks | Cat#QV-4315 |
| Compound 1-3 | Combi-Blocks | Cat#QB-9928 |
| Compound 1-4 | Combi-Blocks | Cat#QH-8016 |
| Compound 1-5 | Requested from NCI | NSC102086 |
| Compound 1-6 | Requested from NCI | NSC131986 |
| Compound 2 | Requested from NCI | NSC63001 |
| Compound 3 | Requested from NCI | NSC204920 |
| Compound 4 | Combi-Blocks | Cat#QZ-0775 |
| Compound 5 | Combi-Blocks | Cat#HA-4857 |
| Compound 6 | Combi-Blocks | Cat#QB-2366 |
| Compound 7 | Ambeed | Cat#A691564 |
| Compound 8 | Ambeed | Cat#A344521 |
| Compound 9 | eNovation Chemicals | Cat#Y4280409 |
| Compound 10 | Ambeed | Cat#A637509 |
| Compound 11 | Ambeed | Cat#A140900 |
| Compound 12 | Ambeed | Cat#A823549 |
| Nicotinamide | Sigma Aldrich | Cat#72340 |
| IP6C | Combi-Blocks | Cat#QJ-2580 |
| Mitomycin C | Apollo scientific (UK) | Cat# BIM0133 |
| **Critical commercial assays** | **Source** | **Identifier** |
| QIAprep Spin Miniprep Kit | QIAGEN | Cat#27104 |
| NEB<sup>®</sup> 10-beta Electrocompetent *E. coli* | New England Biolab | Cat#C3020K |
| BL21(DE3) Competent *E. coli* | New England Biolab | Cat#C2527H |

